# Small RNA directs symbiosis, virulence, and natural products biosynthesis in entomopathogenic bacteria

**DOI:** 10.1101/2020.08.24.265355

**Authors:** Nick Neubacher, Nicholas J. Tobias, Michaela Huber, Xiaofeng Cai, Timo Glatter, Sacha J. Pidot, Timothy P. Stinear, Anna Lena Lütticke, Kai Papenfort, Helge B. Bode

## Abstract

Rapid modulation of gene expression is a key feature for the success of bacteria, particularly for those that rapidly have to adapt to different niches. The lifecycles of *Photorhabdus* and *Xenorhabdus* involve a mutualistic association with nematodes as well as an entomopathogenic phase^1,2^, both of which rely on the production of numerous specialized metabolites (SMs) ^3,4^. Several regulators have been previously implicated in the regulation of SM production in these genera^3,4^. However, the molecular underpinnings regulating SM production and the role of small regulatory RNAs (sRNAs) in this process are unknown. Here we describe the mechanism underlying RNA-mediated control of SM synthesis. We show that the Hfq-dependent sRNA, ArcZ, is an essential requirement for SM production. We discovered that ArcZ directly base-pairs with the mRNA encoding HexA, a key repressor of SM genes. We further demonstrate that the ArcZ regulon is not restricted to SM production, but rather modulates up to ~15% of the transcriptional output in both *Photorhabdus* and *Xenorhabdus*. Together, our study shows that sRNAs are crucial for SM production in these species, reveals previously unknown targets for biosynthetic pathway manipulations, and offers a new tool for the (over)production, isolation and identification of unknown natural products.

---

Regulation via *trans*-encoded sRNAs typically occurs by imperfect base-pairing of sRNAs with their mRNA targets and can be mediated by RNA chaperones such as Hfq and ProQ^13,14^. RNA duplex formation is usually short (6 to 12 nucleotides) and can result in conformational changes in RNA secondary structure with various regulatory outcomes^5^. The RNA chaperone Hfq is highly conserved throughout the bacterial kingdom^6^. Several complex phenotypes have been attributed to Hfq with its regulatory roles being achieved by stabilizing sRNAs and/or mRNAs, mediating base-pairing of sRNAs and their targets, modulation of mRNA translation^6^, as well as accelerating the degradation of sRNAs and their targets^7^. Expression of sRNAs is highly dynamic, with sRNA profiles in *Salmonella* shown to be strongly dependent on the bacterial growth phase^8^. ArcZ is one of the few Hfq-bound sRNAs whose expression remains relatively constant in *Salmonella* throughout the growth phases, making up ~7-12% of all reads identified by Hfq co-immunoprecipitation experiments^8^. ArcZ is transcribed as a 129 nt primary transcript (Figure 1A) and processed into a stable short form (~50 nt) ^8–10^. The processed short form of ArcZ directly activates rpoS translation^10^ and inhibits the expression of several other genes^10^. In *E. coli*, the expression of *arcZ* is repressed by the ArcA-ArcB two-component system under anaerobic conditions. In a negative feedback loop, *arcZ* represses, and is repressed by *arcB* transcription^10^. Although there is a wealth of research on ArcZ in *E. coli* and *Salmonella*^8–10^, its function in other bacteria remains unclear.

**Figure 1A.**
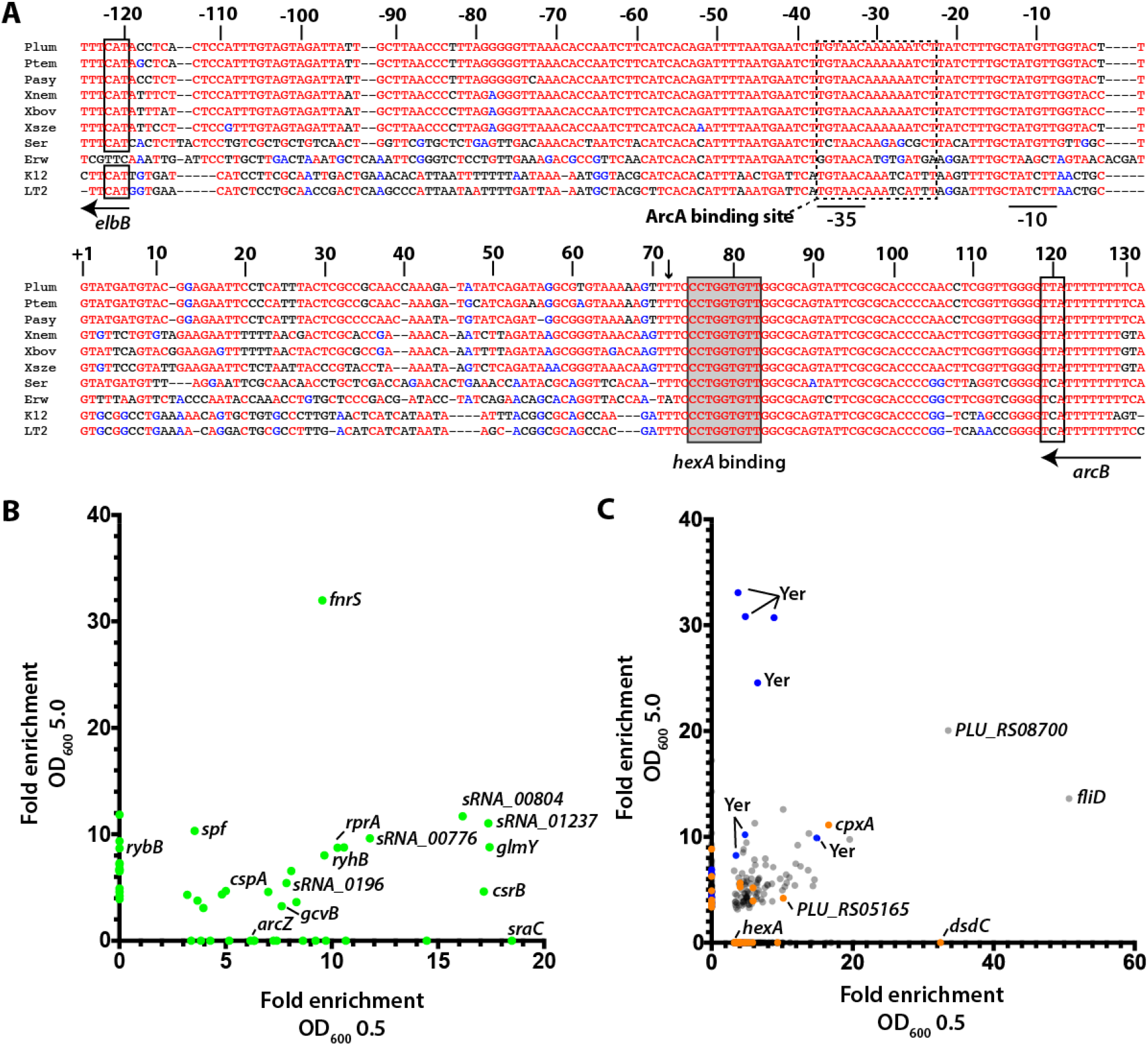
Alignment of *arcZ* sequences from *P. laumondii* TTO1, *P. temperata*, *P. asymbiotica*, *X. nematophila*, *X. bovienii*, *X. szentirmaii*, *Serratia marcescens*, *Erwinia amylovora*, *E. coli* K12 and *Salmonella* typhimurium LT2. Numbers refer to *P. laumondii* sequence. The +1 indicates the transcriptional start of the 129 nt arcZ sequence. Indicated are the start codon of elbB and the stop codon of *arcB*, −10 and −35 binding regions, as well as the conserved ArcA binding region^10^ and the region of base-pairing to *hexA*. The site of ArcZ cleavage is indicated by an arrow. **B** RIPseq enrichment in regions of sRNAs and **C** mRNAs in a strain containing Hfq^3xFLAG^ when compared to the untagged control strain at both optical densities. For a complete list of enriched regions see Supplementary Tables S4 and S5. Blue dots represent SM-related mRNAs, while orange dots represent mRNAs associated with annotated regulators.

SM in bacteria are often responsible for ecologically important activities^11^. In the case of *Xenorhabdus* and *Photorhabdus*, SMs play an essential role in cross-kingdom interactions with nematodes, various insects, as well as bacterial and fungal species competing for the same food source^12^. Our earlier work on *Photorhabdus* showed that deletion of *hfq* resulted in severe perturbation of gene networks, including several key regulators^4^. This led to an overall decrease in SM production and a failure of the bacteria to support their obligate symbiosis with nematodes. Despite SMs playing a central role in the life cycle of the symbiosis, the exact ecological function for many of these compounds remained unknown. Over the past years, significant advances have been made towards finding bioactivities for many of the SMs, with assigned functions including cell-cell communication (photopyrones, dialkylresorcinols^13,14^), nematode development (isopropylstilbene^15^), defense against food-competitors (isopropylstilbene, rhabdopeptides^15,16^), or insect pathogenicity (rhabduscin, rhabdopeptides, glidobactin^16–18^). However, understanding the full potential of SMs in these bacteria is still hampered by a somewhat limited understanding of when individual SMs are produced, and their regulation in general. Regulation of SM in *Photorhabdus* and *Xenorhabdus* so far implicated the regulators Hfq, HexA (also LrhA), LeuO and Lrp^3,4,19,20^. Deletion of *hfq* in *Photorhabdus* resulted in complex regulatory changes, including a strong up-regulation of HexA, a known repressor of SM production^4^. Consequently, SM production was completely abolished in this strain and nematode development was severely restricted.

Given the overlapping lifecycles and niche occupation, we hypothesized that deletion of *hfq* in *Xenorhabdus* would have a similar effect on the production of SMs and the transcriptome. We confirmed this in *X. szentirmaii* DSM16338 using both high-performance LCMS/MS and RNA-seq (Supplementary Results, Supplementary Table S1). To further elucidate the mechanism of SM regulation, we investigated Hfq binding partners. To this end, we sequenced both *X. szentirmaii* and *P. laumondii* using a sRNA sequencing protocol and combined this with CappableSeq data to globally annotate transcriptional start sites belonging to coding sequences or potential novel sRNAs (Supplementary Results, Supplementary Table S2). We confirmed expression of several of these sRNAs by Northern blot analysis (Supplementary Figures S1 & S2). To identify RNA-protein interactions on a global scale, we next employed RNA immunoprecipitation followed by high-throughput sequencing (RIPseq) using chromosomally produced Hfq:3xFLAG protein as bait. We performed these experiments at two different cell densities (*i.e*. OD_600_ 0.5 and OD_600_ 5.0, for a full list of ODs from different experiments, see Supplementary Table S3). From the corresponding sequencing data, we first identified regions of 5 bp or more that were enriched in our tagged Hfq strain (see Methods). We then searched for sRNAs (see Supplementary Results) that were specifically enriched in the tagged samples, when compared to the untagged samples. We identified a total of 37 binding sites in annotated sRNAs (35 unique sRNAs) at OD_600_ 0.5 and 37 binding sites (34 unique) at OD_600_ 5.0 that were enriched by at least three-fold in both replicates (Figure 1B, Supplementary Table S4). During early exponential growth, 11 sRNAs (out of 35) were identified that are described to associate with Hfq in other species, while 10 (out of 34) are known from those that were enriched at OD_600_ 5.0. As a second step, we examined mRNAs enriched in the data. At OD 5.0, 402 mRNAs and 32 annotated 5’-UTRs were identified to associate with Hfq. At OD 0.5 a total of 1,003 mRNAs and 29 5’-UTRs were detected (Figure 1C, Supplementary Table S5).

We hypothesized that the performed Hfq RIP-seq analysis would allow us to identify key sRNAs involved in SM repression. However, our analysis identified >50 potential sRNAs binding Hfq (across both ODs, Figure 1B). Therefore, rather than individually deleting each sRNA, we constructed a transposon mutant library using pSAM-BT_Kan (see Methods and Supplementary Results) and searched for phenotypes consistent with that of the Δ*hfq* strain. The red color afforded to the bacteria by anthraquinone (AQ) production makes the strain especially suitable for transposon mutagenesis when screening for mutants defective in SM biosynthesis. We screened approximately 60,000 clones for obvious phenotypic alterations. Several mutants were defective in some facets of SM production and showed growth defects (Supplementary Results, Supplementary Figure S3, Supplementary Table S6), however, only one displayed the desired phenotype. Re-sequencing of this strain followed by read mapping revealed that the transposon was inserted within an intergenic region associated with the *arcZ* sRNA gene (Supplementary Figure S4).

ArcZ is a well-known Hfq-associated sRNA, which also appeared in our list of Hfq-bound sRNAs in *P. laumondii* (Figure 1B, Supplementary Table S4). To verify that the observed phenotype was derived from the transposon-insertion, we generated a Δ*arcZ* mutant by deleting the major part of the sRNA (Supplementary Figure S4) and a complemented strain by reintroducing an intact version of *arcZ* at the original locus. Northern blot analysis was performed to verify the absence of ArcZ in the deletion mutant and the presence of ArcZ in the WT and the complementation mutant (Figure 2A). RNA sequencing of Δ*arcZ* mutant showed severe transcriptomic changes compared to the WT and Δ*arcZ::arcZ* mutant of *P. laumondii*, reminiscent of that seen in *P. laumondii* Δ*hfq* (Supplementary Results, Supplementary Tables S7 & S8). SM production titers in the Δ*arcZ* strain were strongly decreased, similar to that seen in the transposon-insertion mutant and the complementation strain restored SM production (Figure 2B-H).

**Figure 2A.**
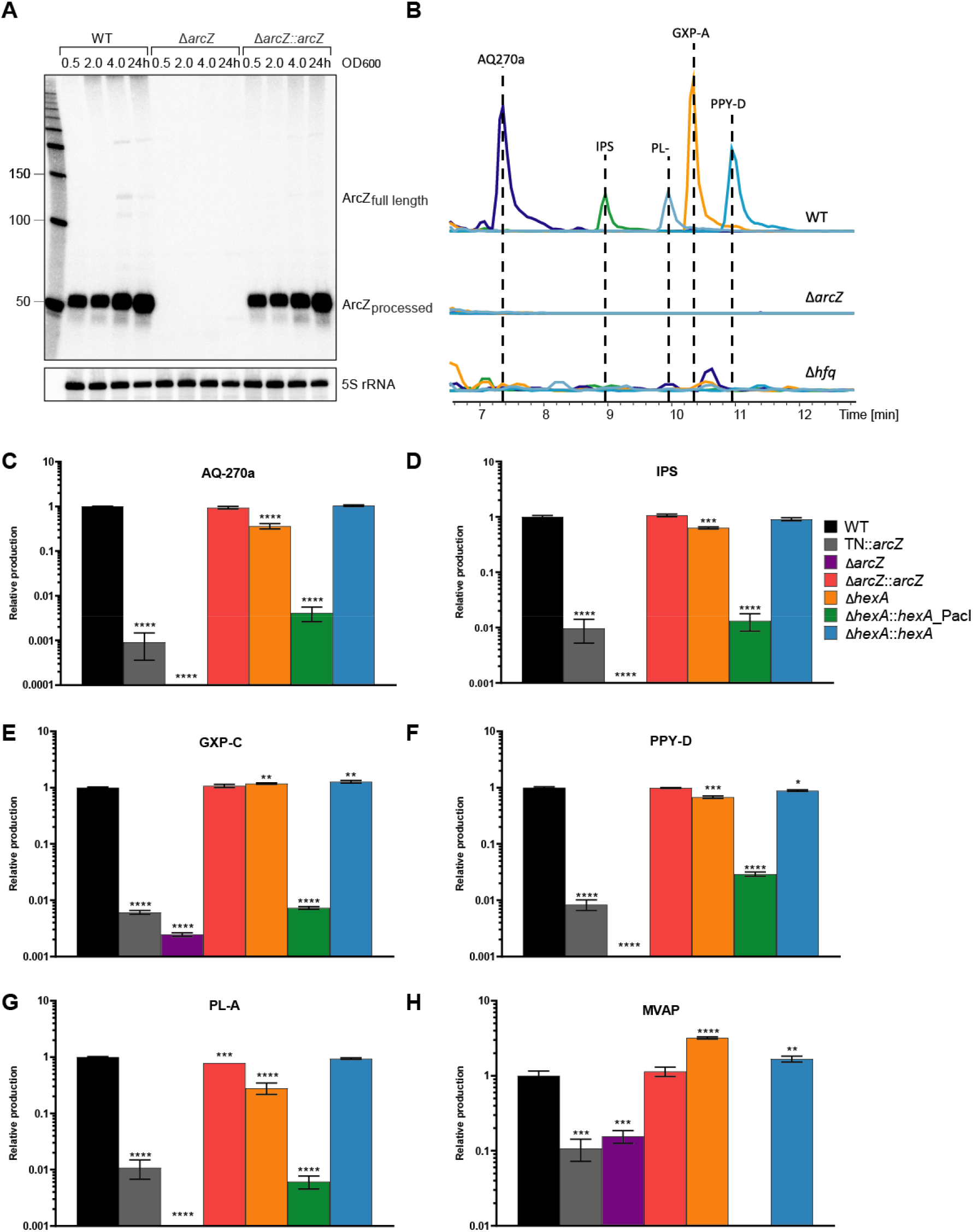
ArcZ expression in *P. laumondii* WT, Δ*arcZ* and Δ*arcZ::arcZ* cells detected by Northern blot analysis. Total RNA samples were collected at three different OD_600_ values (0.5; 2 and 4) and after 24 h of growth. Probing for 5S rRNA served as loading control. **B** Comparison of relative production titers of the major SMs produced by *P. laumondii* WT, Δ*arcZ* and Δ*hfq*. Depicted are the extracted ion chromatograms of anthraquinone (AQ-270a), isopropylstilbene (IPS), phurealipid A (PL-A), GameXPeptide A (GXP-A) and photopyrone D (PPY-D) in the WT and the mutant strains. **C-H**. HPLC-MS quantification of **C**. AQ-270a, **D**. IPS, **E**. GXP-A, **F**. PPY-D, **G**. PL-A and **H**. MVAP in *P. laumondii* WT (black), TN::*arcZ* (grey), Δ*arcZ* (purple), Δ*arcZ::arcZ* (red), Δ*hexA* mutant (orange), Δ*hexA::hexA*_PacI, (green) and Δ*hexA::hexA* (blue). All bars represent relative production in comparison to the wild type. Error bars represent the standard error of the mean. Asterisks indicate statistical significance (* p<0.05, ** p<.005, *** p<0.0005, **** p<0.00005) of relative production compared to WT production levels. Details of all analyzed compounds can be found in Supplementary Table S14.

To corroborate the role of ArcZ in SM production, ArcZ mRNA targets were predicted using CopraRNA^21^ (Supplementary Table S9 & S10). One hit, warranting further investigation was *hexA (lrhA)*, which was previously identified as a highly upregulated gene in our strains and which represses SM production in both *P. laumondii*^22^ and *Xenorhabdus*^3^. CopraRNA predicted a 9 bp-long binding site in the 5’-UTR of *hexA* (Figure 3A). We also identified a corresponding enriched RNA sequence upstream of the *hexA* CDS at OD_600_ 0.5 in the RIPseq experiments (Figure 1C, Supplementary Figure S5). We hypothesized that, through Hfq, ArcZ might bind to the *hexA* transcript leading to repression of HexA. In lab cultures, where SMs are produced, we hypothesized that Hfq and ArcZ prevent HexA production, allowing the strain to synthesize SM. However, if either *hfq* or *arcZ* were deleted, we would expect that *hexA* is no longer repressed, resulting in severely reduced SM production. To test this idea, we altered the predicted site of the ArcZ-*hexA* interaction to a *PacI* restriction site (TTAATTAA) and created a knock-in of *hexA* with the modified sequence in a Δ*hexA* strain (Supplementary Figure S6A&B). We predicted that a knock-in of hexA with an altered 5’-UTR would result in a failure of ArcZ to bind, leading to reduced SM titers. Indeed, the SM production titers in the knock-in mutant with the altered binding site upstream of *hexA* were greatly reduced (Figure 2C-H).

**Figure 3A.**
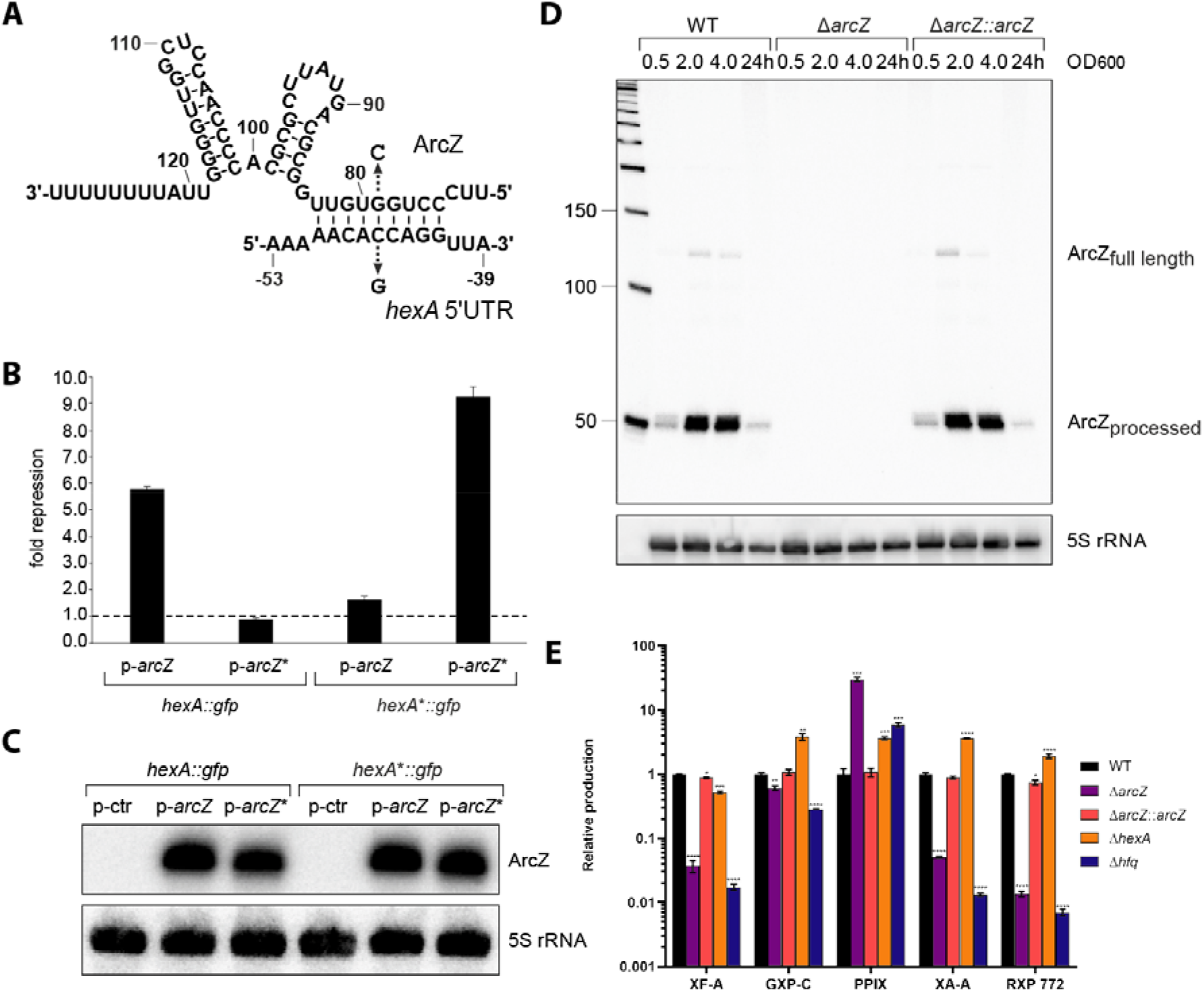
Predicted base-pairing interaction of ArcZ with the 5’-UTR of the *hexA* mRNA. Arrows indicate the single nucleotide mutations tested in B. **B**. Measurement of GFP signals derived from co-expression of a plasmid harbouring the 5’-UTR of *hexA* fused to *gfp* (*hexA::gfp*) or the same fusion with a single point mutation (C-46G, *hexA*\*:*gfp*) with p-ctr, p-*arcZ* or p-*arcZ*\* (G79C). GFP levels of strains carrying the control plasmid (p-ctr) were set to 1. Error bars represent the SD of three biological replicates. **C**. Northern blot analysis of ArcZ expression corresponding to the GFP expression assay shown in B. 5S rRNA was used as loading control. **D**. ArcZ expression in *X. szentirmaii* WT, Δ*arcZ* and Δ*arcZ::arcZ* cells detected by Northern blot analysis. Total RNA samples were collected at three different OD_600_ values (0.5; 2 and 4) and after 24 h of growth. Probing for 5S rRNA served as loading control. **E** HPLC-MS quantification of strains of *X. szentirmaii* WT (grey), Δ*arcZ* mutant (red), Δ*hfq* mutant (orange) complemented arcZ knock-in (green) and Δ*hexA* mutant (blue). All bars represent relative production in comparison to the wild type and were analyzed in triplicate. Shown is the relative production of xenofuranone A (XF-A), GameXPeptide C (GXP-C), protoporphyrin IX (PPIX), xenoamicin A (XA-A) and rhabdopeptide 772 (RXP-772). See also Supplementary Table S6. Error bars represent the standard error of the mean. Asterisks indicate statistical significance (* p<0.05, ** p<0.005, *** p<0.0005, **** p<0.00005) of relative production compared to WT production levels.

To verify the proposed interaction region, we conducted a compensatory base mutation study in *E. coli*. The fifth base-pair of the proposed interaction region was exchanged in the *arcZ* sequence, the *hexA* 5’UTR, or both by site directed mutagenesis (Figure 3A). The *hexA* 5’ UTR sequence was fused to *gfp*. The GFP output was measured to determine the efficiency of inhibition (Figure 3B & C). For the control, the GFP signal derived from the expression of *hexA::gfp* was measured and set to 1. When p-*arcZ* was expressed together with *hexA::gfp*, HexA repression was increased 5.7-fold compared to the control. Additionally, when p-*arcZ*\* (G79C) was expressed, ArcZ* was no longer able to repress HexA. For *hexA*\*::*gfp* (C-46G) in combination with the native ArcZ, HexA repression was only slightly increased compared to the control, suggesting that ArcZ can still bind to the 5’ UTR of hexA but with a much reduced efficiency. When combining p-*arcZ*\* (G79C) with *hexA*\*::*gfp* (C-46G), HexA::GFP repression was increased almost 10-fold, which confirms our hypothesis that ArcZ binds to the 5’-UTR of hexA to repress HexA production. Of note, this base-pairing sequence is located ~50 nts upstream of the hexA translational start site (Fig. 3A) and thus ArcZ binding is unlikely to compete with recognition of the mRNA by 30S ribosomes ^23^. Instead, alignment of the *P. laumondii* hexA 5’ UTR revealed that the ArcZ binding site is CA-rich and highly conserved among other SM-producing bacteria (Supplementary Figure S7). CA-rich sequences located in proximity to translation initiation sites are well-known translational enhancers and sequestration of these regulatory elements by sRNAs has been reported to down-regulate gene expression^24,25^, which might also be relevant for the ArcZ-hexA interaction reported here. In addition, we conducted a proteomic analysis with the WT, Δ*arcZ*, *Δhfq* and *ΔhexA::hexA*_PacI_UTR strains of *P. laumondii*. We used a label free quantification of quadruplicate samples to determine the HexA abundancy in each strain. HexA levels were significantly elevated in all mutant strains (11.8 to 22.7 fold, Supplementary Table 11) compared to the WT, further supporting this mechanism of regulation for SM production.

The *arcZ* gene and its genomic organization are highly conserved among enterobacterial species^9^ (Figure 1A). Since the control of SMs in *Photorhabdus* relays a fundamental ability for these bacteria to occupy their specific niche, we investigated the possibility that the same mechanism occurs in the closely related *Xenorhabdus*. Given the SM reduction in *X. szentirmaii* Δ*hfq*, we constructed a Δ*arcZ* mutant in *X. szentirmaii* in a similar fashion to *P. laumondii*, by deleting 90bp of the predicted *arcZ* sequence. We verified via Northern blots that ArcZ was no longer produced by the deletion mutant and that complementation of the deletion led to production of ArcZ again (Figure 3D). Subsequently, we investigated the transcriptome and SM profile of the WT, deletion and complementation mutant. (Figure 3E, Supplementary Table S12). Consistent with *P. laumondii*, deletion of *arcZ* resulted in a global effect on the transcriptome as well as severely reduced SM titers, both of which was complemented in the Δ*arcZ::arcZ* complementation mutant (Figure 3E, Figure 4).

**Figure 4.**
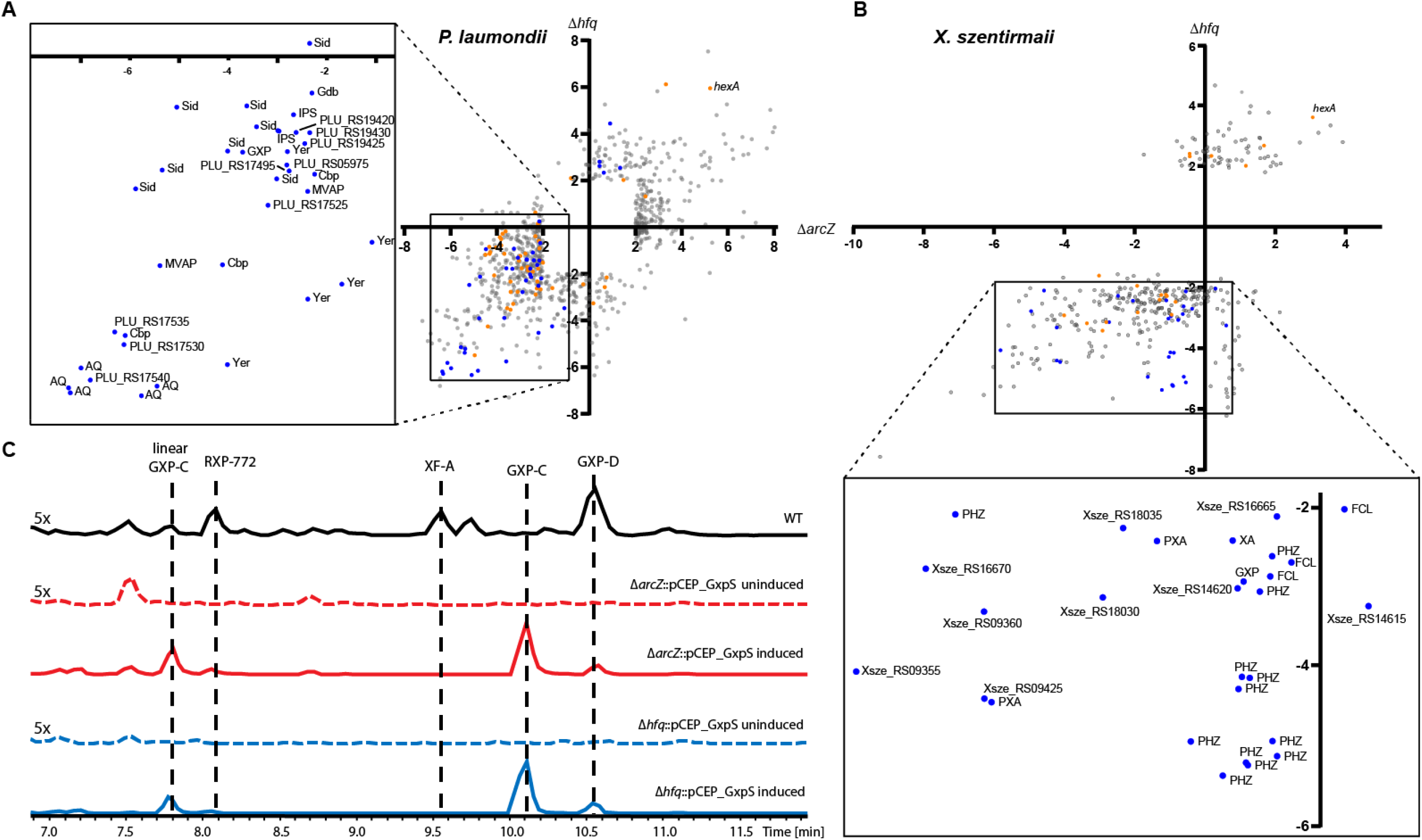
Comparison of ArcZ and Hfq regulon in **A** *P. laumondii* and **B** *X. szentirmaii*. Scatterplots show individual coding sequences and their corresponding regulatory changes compared to wild type in either the Δ*arcZ* (x-axis) or Δ*hfq* (y-axis) mutants, with SMs (blue dots) and regulators (orange dots) highlighted. The inset shows only SM-related coding sequences, including those associated with anthraquinone (AQ), mevalagmapeptide (MVAP), carbapenem (Cbp), yersiniabactin (YER), GameXPeptide (GXP), siderophore (SID), isopropylstilbene (IPS) and glidobactin (Gdb), phenazine (PHZ), fabclavine (FCL), xenoamicin (XA) and pyrrolizixenamide (PXA). **C** Base peak chromatograms (BPCs) of *X. szentirmaii* WT (black), Δ*arcZ*::pCEP_GxpS uninduced (red dotted line), Δ*arcZ*::pCEP_GxpS induced (red solid line), Δ*hfq*::pCEP_GxpS uninduced (blue dotted line) and Δ*hfq*::pCEP_GxpS induced (blue solid line). Peaks corresponding to (cyclo)tetrahydroxybutyrate (THB^53^), Linear GameXPeptide C (GXP-C), rhabdopeptide 772 (RXP), xenofuranone A (XF-A), as well as cyclic GXP-C and GXP-D. Five times zoom was applied to base peak chromatograms in the uninduced and wild type samples.

Our results highlight the critical role of ArcZ in regulating specialized metabolism in these strains. In fact, the critical nature of SM from *Photorhabdus* and *Xenorhabdus* in modulating the insect immune response indicated that ArcZ might be required for niche occupation by these bacteria. In the *P. laumondii* Δ*arcZ* strain, we observed an inability to support nematode development (Supplementary Figure S8), consistent with our earlier observations in the Δ*hfq* mutant^4^. However, the same was not seen in *X. szentirmaii*. We suspect this might be because of the observed increase in protoporphyrin IX (PPIX) production in the *X. szentirmaii* Δ*arcZ* strain. PPIX is a precursor of heme, which is an important cofactor for key biological processes such as oxidative metabolism^26^, protein translation^27^, maintaining protein stability^28^ and many others. However, PPIX cannot be synthesized *de novo* by *Caenorhabditis elegans* and other nematodes^29^. The nematodes therefore rely on external PPIX sources (such as from symbiotic bacteria), which positively affects their growth, reproduction and development^30^. It is interesting that despite *P. laumondii* also being capable of producing PPIX, the *Heterorhabditis* nematode reproduction was not supported in either the Δ*arcZ* mutant, nor the Δ*hfq* strain. This is possibly indicative of the nematode specific requirements for reproduction, which may also include isopropylstilbene as an essential factor in *Heterorhabditis*^15^, where no analogous compound is yet known to be required for *Steinernema*.

In both *Xenorhabdus* and *Photorhabdus*, nearly all analyzed SM-related genes were found to be down-regulated in the Δ*arcZ* mutant, in accordance with the impaired SM production (Figure 4A & B). This provides a chemical background that is devoid of natural products, which allows for isolation and identification of a desired compound due to the absence of compounds with similar retention times. Therefore, Δ*arcZ* mutants could offer a powerful tool for (over-) production and identification of new natural products. As a proof of concept, we conducted a promotor exchange in front of *gxpS* in both *X. szentirmaii* Δ*arcZ* and *X. szentirmaii* Δ*hfq* and compared GXP-C production after induction to the WT (Figure 4C). GXP-C production was found to be increased 90.4 (± 4.7)-fold in *X. szentirmaii* Δ*arcZ*::pCEP_GxpS and increased 138.6 (±17.1)-fold in *X. szentirmaii* Δ*hfq*::pCEP_GxpS compared to the WT (Figure 4C). The striking increase in production, as well as the dramatically reduced chemical background in both strains, highlights the potential of exploiting this regulatory cascade for selective SM production in a strain well-suited for natural product detection. Recently, we showed that this strategy could be applied in a high-throughput manner for rapid screening of bioactivities^31^. The same strategy used here in a Δ*arcZ* strain, demonstrates an alternative route to activation, without the complex perturbations associated with deleting the major RNA chaperone in these bacteria. Interestingly, some comparisons between these mechanisms can be drawn in other SM-producing Enterobacteriaceae (Figure 1A). Erwinia is a genus of plant pathogenic bacteria that produce SMs, where Hfq and ArcZ have both been implicated in virulence^32^, while HexA is a negative regulator of secondary metabolites in these bacteria^33^. Similar parallels can also be seen from *Serratia*^34–36^ and *Pseudomonas*^37^ two other prolific SM producers. Although further investigations will be required to ascertain whether these apparent similarities represent identical mechanisms, the conserved nature of ArcZ in other SM-producing Enterobacteriaceae could suggest that this strategy may yield new avenues for rapid investigation into SM biosynthesis in other taxa.

## Materials & Methods

### Bacterial culture conditions

All *Photorhabdus* and *Xenorhabdus* strains were grown in LB with shaking for at least 16 hours at 30°C. *E. coli* strains were grown in LB with shaking for at least 16 hours at 37°C. The medium was supplemented with chloramphenicol (34 μg/ml), ampicillin (100 μg/ml), rifampicin (50μg/ml) or kanamycin (50 μg/ml) when appropriate. Promotor exchange mutants were induced by adding L-arabinose (2%, v/v) to the cultures. All plasmids and strains used in this study are listed in Supplementary Table S13 & S14.

### Nematode bioassays

All nematodes were cultivated in *Galleria mellonella* and collected on white traps as previously described. Nematode bioassays were also performed as described elsewhere^4^.

### Creation of transposon mutant library

For the transposon mutagenesis, the plasmid pSAM_Kan (containing the mariner transposon) was constructed using pSAM_BT^38^ as a template. To do this, the plasmid was linearized using the primers NN191/NN192. The kanamycin resistance cassette was amplified from the pCOLA_ara_tacI plasmid using the primers NN193/NN194 introducing complementary overhangs to pSAM_BT at both ends of the PCR fragment. The kanamycin resistance cassette was fused with the linearized pSAM_BT plasmid using Hot Fusion cloning thereby replacing the erythromycin resistance cassette with kanamycin resistance. *E. coli* ST18 was transformed with the plasmid pSAM_Kan and further used for the creation of the transposon mutant library of *P. laumondii* TTO1 through conjugation. Transposon-insertion mutants were selected on LB agar containing kanamycin. All primer sequences are listed in Supplementary Table S15.

### Construction of mutant strains

For the deletion of the majority of ArcZ in *P. laumondii* TTO1, a 1123 bp upstream and a 1014 bp downstream product was amplified using the primers NN276/NN277 and NN278/NN279, respectively. The PCR products were fused using the complementary overhangs introduced by the primers and cloned into the PstI and BglII linearized pEB17 plasmid. The resulting plasmid was used for transformation of *E. coli* s17-1 λpir. Conjugation of the plasmid in *P. laumondii* strains and generation of deletion strains by homologous recombination through counter selection was done as previously described ^39^. Deletion mutants were verified by PCR using the primers NN281/NN282 yielding a 632 bp fragment for mutants genetically equal to the WT and a 502 bp fragment for the desired deletion mutant. Complementation of the ArcZ deletion was achieved by inserting the full and intact version of ArcZ at the original locus. To do this, a 2207 bp PCR product was amplified using the primers NN276/NN279 including the upstream and downstream region required for homologous recombination and the full length ArcZ. The fragment was cloned into pEB17 as described above. The verified plasmid construct was used for transformation of *E. coli* s17-1 *λpir* cells. The plasmid was transferred into *P. laumondii* Δ*arcZ* by conjugation and integrated into the genome of *P. laumondii* Δ*arcZ* by homologous recombination. The knock-in mutant was generated by a second homologous recombination through counter selection on LB plates containing 6% sucrose. Knock-in mutants were verified by PCR using the primers NN281/NN282 yielding a 632 bp fragment. The same strategy was used for the construction of the mutant strains in *X. szentirmaii*. To generate the promotor exchange mutants in front of *gxpS*, the plasmid pCEPKMR_ORF00346 was transferred into *X. szentirmaii* Δ*arcZ* and *X. szentirmaii* Δ*hfq* by conjugation and integrated into the genome by homologous recombination.

## DNA extraction

Genomic DNA was extracted using the Gentra Puregene Yeast/Bact Kit (Qiagen) following the manufacturer’s instructions. For sequencing of transposon-insertion mutants, genomic DNA was extracted using the DNeasy Blood & Tissue Kit (Qiagen).

### DNA sequencing and identification of transposon insertion site

DNA isolated from the transposon-insertion mutants was sequenced on the Illumina NextSeq platform. DNA libraries were constructed using the Nextera XT DNA preparation kit (Illumina) and whole genome sequencing was performed using 2 × 150bp paired-end chemistry. A sequencing depth of >50× was targeted for each sample. Genomes were assembled with SPAdes (v 3.10.1) ^40^ and annotated with Prokka v 1.12^41^. Completed genome sequences were analysed and viewed in Geneious v9.1 (https://www.geneious.com).

### RNA extraction, sequencing and analysis

Pre-cultures of *P. laumondii* TTO1, *X. szentirmaii* DSM16338, and their respective ArcZ deletion and knock-in mutants were grown in LB broth overnight with shaking, at 30 °C. The following day, the pre-cultures were used to inoculate fresh LB at an OD_600_ of 0.3. Cells were grown to mid-exponential phase (OD values for each experiment can be found in Supplementary Table S3). RNA was extracted using the RNeasy Mini Kit (Qiagen) following the manufacturer’s instructions. To facilitate cell lysis, the cells were pelleted and snap frozen in liquid nitrogen for 1 min after removing the supernatant. After thawing and resuspending in lysis buffer, the cells were vortexed for 30 sec before proceeding with the protocol. RNA for small RNA libraries were extracted in duplicate, during the mid-exponential phase for *P. laumondii* TTO1 and *X. szentirmaii*.

RNA was sequenced with 150bp paired-end sequencing by Novogene following rRNA depletion with a RiboZero kit and library preparation following the Illumina protocol for strand-specific libraries. Raw data was trimmed using Trimmomatic^42^ and mapped to the reference genome downloaded from NCBI (NC_005126.1 for *P. laumondii* and NZ_NIBV00000000.1 for *X. szentirmaii*) using bowtie2 (v2.3.4.3) ^43^. Resulting .sam files were converted to .bam files using samtools (v1.8) ^44^ and featureCounts (a part of the subread package) ^45^ was used to count reads mapping to annotated genes. Count files were then uploaded to degust (http://degust.erc.monash.edu/) and analyzed using the voom/limma method of normalization. Only genes with an absolute fold change >2 and false discovery rate < 0.01 were considered significantly regulated.

### Northern blot analysis

For Northern blot analysis, total RNA was prepared and analyzed as described previously^46^. Briefly, RNA samples were separated on 6% polyacrylamide / 7 M urea gels and transferred to Hybond–XL membranes (GE Healthcare) by electro-blotting. Membranes were hybridized in Roti-Hybri-Quick buffer (Roth) at 42°C with gene-specific [^32^P] end-labeled DNA oligonucleotides, and washed in three subsequent steps with SSC (5x, 1x, 0.5x) / 0.1% SDS wash buffer. Signals were visualized on a Typhoon FLA 7000 phosphorimager (FUJIFILM). Oligonucleotides for Northern blot analyses are listed in Supplementary Table S15.

### Compensatory base mutation and GFP fluorescence assay

Plasmids pMH078 and pMH079 were generated using Gibson assembly^47^. For plasmid pMH078 the *arcZ* gene was amplified using *P. laumondii* TT01 genomic DNA with oligonucleotides KPO-6147 and KPO-6148 and fused into a pEVS143 vector backbone^48^, linearized with KPO-0092 and KPO-1397. To construct plasmid pMH079, the 5’UTR and the first 20 aa of *hexA* were amplified using *P. laumondii* TT01 genomic DNA with KPO-6145 and KPO-6146, and the pXG10-gfp vector^49^ was linearized with KPO-1702 and KPO-1703. pMH078 and pMH079 served as templates to insert single point mutations in the *arcZ* gene as well as the *hexA* 5’UTR using site-directed mutagenesis and oligonucleotide combinations KPO-6156/KPO-6157 and KPO-6164/KPO-6165, respectively, yielding plasmids pMH080 and pMH081.

Target regulation using GFP reporter fusions was analyzed as described previously^49^. *E. coli* Top10 cells were grown overnight in LB medium (37°C, 200 rpm shaking conditions). Three independent cultures were used for each strain. Cells were washed in PBS and GFP fluorescence intensity was determined using a Spark 10 M plate reader (TECAN). Control samples not expressing fluorescence proteins were used to subtract background fluorescence.

### CappableSeq analysis

Cappable seq was performed as previously described^50^ by Vertis Biotechnologies (Germany). Raw sequences were trimmed with Trimmomatic^42^ and mapped with bowtie2^43^ to NC_005126.1 for *P. laumondii* and NZ_NIBV00000000.1 for *X. szentirmaii*. Transcriptional start sites were detected using readXplorer’s (v2.2.3) ^51^ built in TSS detection function with the following settings: use only single perfect matches, minimum number of read starts = 100, minimum percent coverage increase = 750, detect novel transcripts, min. transcript extension = 40, max distance to feature of leaderless transcripts = 5, associate neighbouring TSS within 3bp.

### RIP-seq analysis

Overnight cultures of *P. laumondii* TTO1 (WT and Hfq^3xFLAG^) were inoculated into fresh LB media in duplicate and grown at 30 °C with shaking at 200 rpm. Bacteria were harvested by centrifugation at 4°C, 4000 rpm for 15 min when cells reached OD_600_=0.5 and OD_600_=5.0. Cell pellets were resuspended in 1 mL lysis buffer (20 mM Tris pH 8.0, 150 mM KCl, 1 mM MgCl2, 1 mM DTT) and pelleted again by centrifugation (5 min, 11,200 g, 4°C). The supernatants were discarded and the pellets were snap-frozen in liquid nitrogen. After thawing on ice, cells were resuspended in 800 μl lysis buffer and transferred into tubes containing 300 μl glass beads to break cells via a Bead Ruptor (150 sec, twice, 2 min break on ice in between). After short centrifugation (15,000 g, 4°C), lysates were transferred into fresh precooled tubes and centrifuged for 30 minutes at 15,200 g at 4°C. The cleared lysates were transferred into new tubes and incubated with 35 μl FLAG-antibody (Monoclonal ANTI-FLAG M2, Sigma, #F1804) with rotation for 45 min at 4°C, followed by addition of 75 μl Protein G Sepharose (Sigma, #P3296) and rotating for 45 min at 4°C again. After five wash steps with lysis buffer (via inverting the tube gently and centrifuging for 4 min at 4° C), samples were subjected to RNA and protein separation by Phenol:Chloroform:Isoamylalcohol (P:C:I; 25:24:1, pH 4.5, Roth) extraction. The upper phase (~ 500 μl) was transferred into to a new tube and precipitated overnight at −20°C with 1.5 ml EtOH:Na(acetate) (30:1) and 1.5 μl GlycoBlue (#AM9516, Ambion). After centrifugation for 30 minutes at 11,200 rpm at 4°C, RNA pellets were washed with 500 μl 70% EtOH, dried and resuspended in 15.5 μl nuclease-free H_2_O. RNA was treated with 2 μl DNase I, 0.5 μl RNase inhibitor and 2 μl 10× DNase buffer at 37°C for 30 min. Afterwards, samples were supplemented with 100 μl H_2_O, and again subjected to P:C:I extraction. The upper phase (~120 μl) was transferred into a new tube with addition of 2.5-3 volumes (~350 μl) of EtOH:Na(acetate) (30:1) and stored at −20°C overnight for RNA precipitation. RNA pellets were harvested via centrifugation for 30 min at 13,000 rpm, 4 °C, and washed with 500 μl 70% EtOH, dried and resuspended in nuclease-free H_2_O. cDNA libraries were prepared using the NEBNext Small RNA Library Prep Set for Illumina (NEB, #E7300S) according to the manufacturer’s instructions and sequenced on a HiSeq 1500 system in single-read mode with 100 nt read length.

For RIPseq analysis, the enriched/control sample pairs were normalized by the number of raw reads present after trimming. Depth counts of all samples were obtained using samtools (v1.8)^44^. Only nucleotide positions with a depth of at least 50 reads in the enriched samples were taken for further analysis. The corresponding depth in the unenriched samples was matched for each nucleotide. A region was considered to be enriched if the enrichment factor was at least three and the corresponding ‘enriched’ nucleotide was present in both sample pairs. Finally, we considered a region to be enriched if more than five consecutive nucleotides were identified as enriched.

### ArcZ binding prediction

ArcZ from *E. coli* was used to define the boundaries of ArcZ in *Xenorhabdus* and *Photorhabdus*. We then took our annotated ArcZ sequence together with several ArcZ homologs from other Enterobacteriaceae (listed in Supplementary Table S9) and used the online CopraRNA tool^21^, a part of the Freiburg RNA tools suite^52^, with default parameters.

### Metabolite extraction and HPLC-MS/MS analysis

Fresh 10 ml of LB was inoculated with an overnight culture to an OD_600_ = 0.1. After 72 h of cultivation at 30°C with shaking, 1 ml of the culture was removed from the culture, centrifuged for 20 min at 13,300 rpm and the supernatant was directly subjected for HPLC-UV/MS analysis using a Dionex Ultimate 3000 system with a Bruker AmaZon X mass spectrometer. The compounds peak areas were quantified using TargetAnalysis 1.3 (Bruker). All analyzed compounds are listed in Supplementary Table S16.

### Proteome analysis

The detail of the proteomics procedure can be assessed in (https://pubs.acs.org/doi/abs/10.1021/acs.jproteome.8b00216). In short, to extract proteins from E.coli, frozen cell pellets 300 μL lysis buffer (0.5% Na-desoxycholate in 100 mM NH_4_HCO_3_) were added to the cell pellet, and incubated at 95°C for 10. The protein concentration in the supernatant was determined with a BCA Protein Assay Kit (Thermo Fisher, #23252). Reduction and alkylation was performed at 95 °C using 5mM TCEP and 10mM Chloroacetamide for 15min. 50 μg of protein was transferred to new reaction tubes and protein digestion was carried out overnight at 30 °C with 1 μg trypsin (Promega). After digest, the peptides were desalted using CHROMABOND Spincolumns (Macherey-Nagel) that were conditioned with 500 μL of acetonitrile and equilibrated with 500 μL and 150 μL 0.1% TFA. After loading the peptides were washed with 500 μL 0.1% TFA in 5:95 acetonitrile:water, peptides were eluted with 400 μL 0.1% TFA in 50:50 acetonitrile:water. Peptides were concentrated and dried under vacuum at 50°C and dissolved in 100 μL 0.1% TFA by 25 s of sonication and incubation at 22°C under shaking at 1200 rpm for 5 min. 1 μg peptide was analyzed using liquid chromatography-mass spectrometry (LC-MS/MS).

The LC-MS/MS analysis including lable-free quantification was carried out as previously described in (https://pubs.acs.org/doi/abs/10.1021/acs.jproteome.8b00216) with minor modifications.

LC-MS/MS analysis of protein digests was performed on Q-Exactive Plus mass spectrometer connected to an electrospray ion source (Thermo Fisher Scientific). Peptide separation was carried out using Ultimate 3000 nanoLC-system (Thermo Fisher Scientific), equipped with packed in-house C18 resin column (Magic C18 AQ 2.4 μm, Dr. Maisch). The peptides were first loaded onto a C18 precolumn (preconcentration set-up) and then eluted in backflush mode with a gradient from 98 % solvent A (0.15 % formic acid) and 2 % solvent B (99.85 % acetonitrile, 0.15 % formic acid) to 35 % solvent B over 30 min. Label-free quantification was done using Progenesis QI software (Nonlinear Dynamics, v2.0), MS/MS search was performed in MASCOT (v2.5, Matrix Science) against the Uniprot *Photorhabdus* laumondii protein database. The following search parameters were used: full tryptic search with two missed cleavage sites, 10ppm MS1 and 0.02 Da fragment ion tolerance. Carbamidomethylation (C) as fixed, oxidation (M) and deamidation (N,Q) as variable modification. Progenesis outputs were further processed with SafeQuant (https://www.ncbi.nlm.nih.gov/pubmed/23017020).

## Data availability

All .mzXML files from HPLC-MS runs are available at MassIVE (https://massive.ucsd.edu) under the ID MSV000084163. Raw sequence data is available at the European nucleotide archive under project accession numbers PRJEB33827 and PRJEB24159. Updated annotation files are available in .gff format under Supplementary Tables S17 (*P. laumondii*) and S18 (*X. szentirmaii*). The proteomic data can be accessed at PRIDE (https://www.ebi.ac.uk/pride/) with the project accession number PXD019095.

## Acknowledgements

This work was funded in part by the DFG (SFB 902, Project B17) and the LOEWE Centre for Translational Biodiversity Genomics (TBG) supported by the State of Hesse. K.P. acknowledges funding by DFG (EXC 2051, Project 390713860), the Vallee Foundation, and the European Research Council (StG-758212). We thank Andrew Goodman for providing pSAM-BT and helpful discussions. We thank Laura Pöschel and Antje K. Heinrich for plasmid construction.

