## Supplementary Information for "Small RNA directs symbiosis, virulence, and natural products biosynthesis in entomopathogenic bacteria"

**Supplementary Table S1.** Significantly affected genes in *X. szentirmaii* (∆*hfq* RNASeq)

**Supplementary Table S2.** Transcriptional start sites identified for *X. szentirmaii*

**Supplementary Table S3.** OD_600_ values of cultures from all experiments

**Supplementary Table S4.** Enriched sRNA sequences from RIPseq experiment

**Supplementary Table S5.** Enriched mRNA and 5’-UTR sequences from RIPseq experiment

**Supplementary Table S6.** OD_600_ of *P. laumondii* WT and TN-mutants after 72 h of cultivation.

**Supplementary Table S7.** Significantly affected genes in TT01 ∆*hfq* and ∆*arcZ*

**Supplementary Table S8.** Significantly affected genes in TT01 only ∆*arcZ*

**Supplementary Table S9.** ArcZ sequences used in Copra analysis

**Supplementary Table S10.** Top 200 hits defined by Copra as potentially interacting with ArcZ

**Supplementary Table S11.** Proteomic analysis of *P. laumondii* WT, ∆*arcZ*, ∆*hfq* and ∆*hexA*::*hexA*_*Pac*I_UTR

**Supplementary Table S12.** Significantly affected genes in *X. szentirmaii* (∆*arcZ* RNASeq)

**Supplementary Table S13.** Plasmids used in this study

**Supplementary Table S14.** Strains used in this study

**Supplementary Table S15.** Primer sequences used in this study

**Supplementary Table S16.** Compounds targets used in HPLC-MS analyses

**Supplementary Table S17.** Updated *P. laumondii* TTO1 annotation formatted as .gff

**Supplementary Table S18.** Updated *X. szentirmaii* DSM16338 annotation formatted as .gff

**Supplementary Figure S1.** Expression of various sRNAs in *P. laumondii* at different time points

**Supplementary Figure S2.** Expression of various sRNAs in *X. szentirmaii* at different time points.

**Supplementary Figure S3.** Phenotype of transposon insertion mutants of *P. laumondii*

**Supplementary Figure S4.** Sequence of region in *P. laumondii* TTO1 containing predicted *arcZ* sequence

**Supplementary Figure S5.** RIPseq enrichment around the region of *hexA*

**Supplementary Figure S6.** 5’ UTR of *hexA* including predicted ArcZ binding site

**Supplementary Figure S7.** Alignment of the *hexA* 5’ UTR from different species

**Supplementary Figure S8.** Nematode development assays

**Supplementary Figure S9.** The ArcZ and Hfq regulon in *P. laumondii*

**Supplementary Results**

**Hfq is involved in SM biosynthesis in *Xenorhabdus.*** To confirm if Hfq is also involved in SM regulation in *Xenorhabdus*, we created a knockout of *hfq* in *X. szentirmaii* DSM16338 and performed HPLCMS/MS and RNAseq on the confirmed deletion strains (Supplementary Table S1). In contrast to *Photorhabdus*, the *X. szentirmaii* ∆*hfq* strain only revealed 312 coding sequences significantly regulated compared to the wild type at mid-exponential phase. In accordance with our hypothesis, *hexA* was significantly upregulated (8.4x, FDR<0.01, Supplementary Table S1). Consistent with this observation, the production of nearly all known SMs were decreased (Figure 3E), suggesting a conserved mode of action in *Xenorhabdus*.

**Identification of sRNAs in *Photorhabdus* and *Xenorhabdus.*** Only very little is known about sRNAs from entomopathogenic bacteria. To identify potential Hfq-binding sRNAs and consequently the Hfq-based regulation of SMs in general, we sequenced the RNA of *P. laumondii* (formerly *P. luminescens*) and *X. szentirmaii* using a library preparation protocol specific for sRNAs. Sequences of the sRNAs from two libraries from each of *Photorhabdus* and *Xenorhabdus* yielded a total of 26,784,563 (13,204,857 and 13,579,706) and 28,813,442 (13,472,683 and 15,340,759) raw reads, respectively. Additionally, we prepared samples from *X. szentirmaii* for CappableSeq, a protocol that differentiates between primary and secondary transcripts ^1^. We recently reported a data set from *P. laumondii*, which identified 15,500 primary and 3,741 secondary transcripts ^2^. Here, we reanalyzed these data using stricter cutoff criteria (see Methods) resulting in a total of 6,174 TSSs. The *X. szentirmaii* CappableSeq data led to the identification of 2,196 TSSs (Supplementary Table S2).

By combining data from the CappableSeq experiments data along with RNAseq data from ∆*hfq* and wild type strains (also ∆*hfq*∆*hexA* and ∆*hfq*::*hfq* in *Photorhabdus* from our previous study^3^), we were able to annotate putative transcripts, 5’-untranslated regions (UTRs), 3’-UTRs and sRNAs using ANNOgesic^4^ (Supplementary Table S17 and S18). The annotated sRNAs were added to those described in the Bacterial sRNA Database (BSRD)^5^ yielding a total of 280 and 130 candidates for sRNAs in *Photorhabdus* and *Xenorhabdus*, respectively (Supplementary Table S17 & S18).

**Transposon mutant library screen.** A transposon mutant library was constructed to identify genes defective in SM production. Many of the analysed mutant strains showed severely reduced SM production titers in comparison to the WT strain. In most cases, multiple SM classes were affected by the transposon insertion (Supplementary Figure S3). On rare occasions, the transposon insertion led to an increase in production of certain SMs. For example, dmPLA-A and MVAP levels were elevated in mutant strain 9 and IPS titers were slightly raised in the TN-mutant strains 10 and 11. Interestingly, the remaining SMs were negatively affected in those strains. As the growth appeared to be affected by the transposon insertion (Supplementary Table S6, it remains uncertain how the growth defects correlate with SM production. For further analysis, we decided to focus on strain 3 that showed only moderate growth defects while at the same time producing reduced SM titers, consistent with the phenotype of the *hfq* deletion mutant.

**The ArcZ regulon in *Photorhabdus* and *Xenorhabdus.*** Since there is a clear overlap between the regulons and functions of Hfq and ArcZ, we performed RNAseq on the ∆*arcZ* strains of *P. laumondii* and *X. szentirmaii*, as well as on their respective knock-in complementation mutants. RNAseq analysis on the deletion of *arcZ* in *Photorhabdus* revealed an even broader effect than in our ∆*hfq* mutant, significantly affecting the transcriptional level of 735 coding sequences in *P. laumondii* (FDR<0.01; log_2_ fold change >2, Figure 4A, Supplementary Table S7 & S8). In *X. szentirmaii*, a global effect of the *arcZ* deletion was also observed, albeit only 191 genes were affected in this strain (Supplementary Table S12). In both deletion strains however, the majority of affected coding sequences were downregulated (Figure 4A & B, Supplementary Table S8 & S12). In an attempt to identify broader effects, we grouped all the genes that were significantly changed into eight different categories based on their known or proposed function: SM, regulators, virulence, phage related, cell wall, cell processes, hypothetical proteins and unknown. We first included only those genes that were significantly regulated in the *arcZ* deletion mutant and not in the *hfq* deletion mutant (Supplementary Figure S9A). In all cases (except for virulence related and unknown) a clear trend towards downregulation of the transcriptional level could be observed in the deletion of *arcZ*. This trend was also observed in the *hfq* deletion mutant, although somewhat weakened compared to the *arcZ* deletion strain. The knock-in complementation restored the vast majority of observed changes back to WT level (Supplementary Figure S9A). Finally, we looked at genes whose expression was significantly altered in both the *arcZ* and *hfq* deletion strain. The individual categories clustered very closely together as indicated by the median (Supplementary Figure 9B).

**Effect of *arcZ* deletion in *Xenorhabdus.*** The drastic reduction in SMs in the deletion mutant was restored with a knock-in complementation of *arcZ* (Supplementary Figure S3). We also observed that protoporphyrin IX (PPIX), the direct precursor for heme, was highly overproduced (~30-fold) in the ∆*arcZ* strain of *X. szentirmaii* compared to the WT, suggesting that the regulatory functions of ArcZ in *Photorhabdus* and *Xenorhabdus* possibly go beyond SM production. Since heme is reported to play an important role in nematode growth and development, we used the deletion mutants and complemented strains and performed nematode development assays. Both the WT and ∆*arcZ* strain of *X. szentirmaii* were able to support nematode development after 4 days of inoculation. However, the ∆*arcZ* strain of *P. laumondii* showed a significantly reduced capability to support nematode development (Supplementary Figure S8), consistent with our data showing that isopropylstilbene falls under the Hfq-ArcZ regulatory umbrella (Figure 4A, Supplementary Tables S7 & S8).

**Supplementary Table S3**. Optical density values for sequencing experiments

| **Species** | **OD_600_ value** | **Purpose** |
| --- | --- | --- |
| *P. laumondii* replicate A | 5.8 | sRNA sequencing |
| *P. laumondii* replicate B | 5.4 | sRNA sequencing |
| *X. szentirmaii* replicate A | 5.5 | sRNA sequencing |
| *X. szentirmaii* replicate B | 6.1 | sRNA sequencing |
| *P. laumondii* replicate A | 4 | RNA-sequencing |
| *P. laumondii* replicate B | 4.3 | RNA-sequencing |
| *P. laumondii* ∆*arcZ* replicate A | 3.8 | RNA-sequencing |
| *P. laumondii* ∆*arcZ* replicate B | 4.1 | RNA-sequencing |
| *P. laumondii* knock-in replicate A | 5.2 | RNA-sequencing |
| *P. laumondii* knock-in replicate B | 4.6 | RNA-sequencing |
| *X. szentirmaii* replicate A | 4.3 | RNA-sequencing (control for ∆*arcZ* mutant) |
| *X. szentirmaii* replicate B | 3.8 | RNA-sequencing (control for ∆*arcZ* mutant) |
| *X. szentirmaii* replicate A | 4.5 | RNA-sequencing (control for ∆*hfq* mutant) |
| *X. szentirmaii* replicate B | 4.5 | RNA-sequencing (control for ∆*hfq* mutant) |
| *X. szentirmaii* ∆*arcZ* replicate A | 4.8 | RNA-sequencing |
| *X. szentirmaii* ∆*arcZ* replicate B | 4.3 | RNA-sequencing |
| *X. szentirmaii* ∆*hfq* replicate A | 4.8 | RNA-sequencing |
| *X. szentirmaii* ∆*hfq* replicate B | 4.7 | RNA-sequencing |
| *X. szentirmaii* knock-in replicate A | 3.6 | RNA-sequencing |
| *X. szentirmaii* knock-in replicate B | 3.8 | RNA-sequencing |
| *P. laumondii* replicate A | 0.5 | RIP-Seq |
| *P. laumondii* replicate B | 0.5 | RIP-Seq |
| *P. laumondii* replicate A | 5 | RIP-Seq |
| *P. laumondii* replicate B | 5 | RIP-Seq |

**Supplementary Table S6.** OD_600_ of *P. laumondii* WT and TN-mutants after 72 h of cultivation.

| **Strain** | **OD_600_** |
| --- | --- |
| WT | 14.5 |
| 1 | 9.8 |
| 2 | 8.9 |
| 3 | 7.6 |
| 4 | 1.7 |
| 5 | 3.9 |
| 6 | 12.5 |
| 7 | 3.5 |
| 8 | 14.8 |
| 9 | 5.4 |
| 10 | 9.2 |
| 11 | 4.8 |

**Supplementary Table S9**. ArcZ sequences used for CopraRNA Analysis

| **Species** | **Genome accession** | **Sequence (5’-3’)** |
| --- | --- | --- |
| *Escherichia coli* K12 | NC_000913 | gugcggccugaaaaacagugcugugcccuuguaacucaucauaauaauuuacggcgcagccaagauuucccugguguuggcgcaguauucgcgcaccccggucuagccggggucauuuuuu |
| *Citrobacter koseri* ATCC BAA-895 | NC_009792 | gugcggccugaaaagcagagcugcgccuguguaaaaaacaaucauaacuuacggcgcagccacgauuucccugguguuggcgcaguauucgcgcaccccggucaauccggggucauuuuuu |
| *Citrobacter rodentium* ICC168 | NC_013716 | gugcggccugaaaaugagcgcugcgcguuuaaaauaugagaauaacuuaccgcgcagcuacgauuucccugguguuggcgcaguauucgcgcaccccgguuaauccggggucauuuuuu |
| *Escherichia fergusonii* ATCC 35469 | NC_011740 | gugcggccugaaaaacagugcugcgccuugguuacaaacgacaauaauuuacggcgcagccauaauuucccugguguuggcgcaguauucgcgcaccccgguuaauccggggucauuuuuu |
| *Pectobacterium carotovorum* subsp. *carotovorum* PC1 | NC_012917 | uuaagacacgaauauccgcacuugcgaguuuacaaaaccugaaaucuaaaugcaggugcauguuuucccugguguuggcgcauaauucgcgcaccccggcuucggccggggucauuuuuu |
| *Salmonella enterica* subsp. *enterica* serovar Typhimurium str. LT2 | NC_003197 | gugcggccugaaaacaggacugcgccuuugacaucaucauaauaagcacggcgcagccacgauuucccugguguuggcgcaguauucgcgcaccccggucaaaccggggucauuuuuu |
| *Serratia proteamaculans* 568 | NC_009832 | cuguaugauguuuaggaauucucuacaaccugcucgaccagaacauuaaaaccaauacgcagguucacaauuucccugguguuggcgcaauauucgcgcaccccggccuaggucggggucauuuuuu |
| *Yersinia pestis* CO92 | NC_003143 | guaugauguaugaaagaauccugacaaccugcgaauucacucgaaaucgaaauaauacgcagguuaacguuuucccugguguuggcgcagucuucgcgcaccccggccucggucgggguuauuuuuu |

**Supplementary Table S13**. Plasmids used in this study

| **Plasmid** | **Genotype** | **Reference** |
| --- | --- | --- |
| pSAM_BT | R6K ori, Amp^R^, oriT, Himar1C9 transposase, transposon containing *ermG* | ^6^ |
| pSAM_KAN | R6K ori, Amp^R^, oriT, Himar1C9 transposase, mariner transposon containing Kan^R^ | This study |
| pCK_cipB | R6K ori, CM^R^, oriT, *sacB*, *traI* | ^7^ |
| pCOLA_ara_tacI | cola oriR, Kan^R^ | ^8^ |
| pCKcipB_∆*arcZ*_TT01 | R6K ori, CM^R^, oriT, *sacB*, *traI*, containing a 1083 bp upstream and a 994 bp downstream region of *arcZ* | This study |
| pCKcipB_∆*arcZ*_XSZ_DSM | R6K ori, CM^R^, oriT, *sacB*, *traI*, containing a 860 bp upstream 965 bp downstream region of *arcZ* | This study |
| pCKcipB_hfq_flag_TT01 | R6K ori, CM^R^, oriT, *sacB*, *traI*, containing the upstream and downstream region of *hfq* including a 3xFLAG tag that was introduced via primers | This study |
| pEB17 | R6K ori, Kan^R^, oriT, *sacB*, *traI* | This study |
| pEB17_*arcZ*_TT01 | R6K ori, Kan^R^, oriT, *sacB*, *traI*, containing a 2207 bp fragment including a 1083 bp upstream and a 994 bp downstream region of *arcZ* and the full version of *arcZ* | This study |
| pEB17_*arcZ*_XSZ_DSM | R6K ori, Kan^R^, oriT, *sacB*, *traI,* containing a 1915 bp fragment including a 860 bp upstream and a 965 bp downstream region of *arcZ* and the full version of *arcZ* | This study |
| pEB17_∆*hexA*_XSZ | R6K ori, Kan^R^, oriT, *sacB*, *traI*, containing a 1110 bp upstream 1036 bp downstream region of *hexA* | This study |
| pEB17_*HexA*_*Pac*I_TT01 | R6K ori, Kan^R^, oriT, *sacB*, *traI*, containing a 2141 bp fragment (includes upstream region of *hexA*, the complete *hexA* gene and downstream region of *hexA* with the introduced *Pac*I restriction site instead of the predicted ArcZ binding site) | This study |
| pCEPKMR_ORF00346 | R6K ori, Kan^R^, oriT, *traI*, containing a 618 bp fragment homologous to *gxpS* required for homologous recombination | This study |
| pMH078 | p-*arcZ*, constitutive sRNA expression plasmid, P15A, Kan^R^ | This study |
| pMH079 | pXG10-*hexA::gfp*, containing the 5’UTR and the first 20 aa of *hexA*, pSC101*, Cm^R^ | This study |
| pMH080 | p-*arcZ** (G79C), constitutive sRNA expression plasmid, P15A, Kan^R^ | This study |
| pMH081 | pXG10-*hexA*::gfp* (C-46G)*,* containing the 5’UTR and the first 20 aa of *hexA*, pSC101*, Cm^R^ | This study |
| p-ctr | pCMW-1, control plasmid, P15A, Kan^R^ | ^9^ |

| **Supplementary Table S14**. Bacterial strains used in this study | | |
| --- | --- | --- |
| **Strain** | **Genotype/Description** | **Reference** |
| ***Escherichia coli* S17-1 λpir** | Tp^R^ Sm^R^ *recA*, *thi*, *pro*, *hsdR*-M+RP4: 2- Tc:Mu:Km Tn7 λ*pir* | Invitrogen |
| S17-1 λpir + pCKcipB_∆*arcZ*_TT01 | S17-1 λ*pir* + pCKcipB_∆*arcZ*_TT01, Cm^R^, R6K ori | This study |
| S17-1 λpir + pCKcipB_∆*arcZ*_XSZ_DSM | S17-1 λ*pir* + pCKcipB_∆*arcZ*_XSZ_DSM, Cm^R^, R6K ori | This study |
| S17-1 λpir + pEB17_ *arcZ*_TT01 | S17-1 λ*pir* + pEB17_*arcZ*_TT01, Kan^R^, R6K ori | This study |
| S17-1 λpir + pEB17_ *arcZ*_XSZ_DSM | S17-1 λ*pir* + pEB17_*arcZ*_XSZ_DSM, Kan^R^, R6K ori | This study |
| S17-1 λpir + pEB17_ ∆*hexA*_XSZ_DSM | S17-1 λ*pir* + pEB17_∆*hexA*_XSZ_DSM, Kan^R^, R6K ori | This study |
| S17-1 λpir + pEB17_ *hexA_*PacI_TT01 | S17-1 λ*pir* + pEB17*_hexA_*PacI_TT01, Kan^R^, R6K ori | This study |
| S17-1 λpir + pCEPKMR_ORF00346 | S17-1 λ*pir* + pCEPKMR_ORF00346, Kan^R^, R6K ori | This study |
| ***E. coli* ST18** | S17 λpir ∆*hemA* | ^10^ |
| ST18 + pCKcipB_hfq_flag_TT01 | ST18 + pCKcipB_hfq_flag_TT01 | This study |
| ***E. coli* Top10** | *F- mcrA Δ(mrr-hsdRMS-mcrBC) φ80lacZΔM15 ΔlacX74 nupG recA1 araD139 Δ(ara-leu)7697 galE15 galK16 rpsL(StrR) endA1 λ-* | Invitrogen |
| Top10 + pMH079 + p-ctr | Top10 + pXG10-*hexA*::gfp+ p-ctr | This study |
| Top10 + pMH079 + pMH078 | Top10 + pXG10-*hexA*::gfp+ p-*arcZ* | This study |
| Top10 + pMH079 + pMH080 | Top10 + pXG10-*hexA*::gfp+ p-*arcZ** | This study |
| Top10 + pMH081 + p-ctr | Top10 + pXG10-*hexA**::gfp+ p-ctr | This study |
| Top10 + pMH081 + pMH078 | Top10 + pXG10-*hexA**::gfp+ p-*arcZ* | This study |
| Top10 + pMH081 + pMH080 | Top10 + pXG10-*hexA**::gfp+ p-*arcZ** | This study |
| ***Photorhabdus laumondii subsp. laumondii* TT01** | WT, Rif^R^ | ^11^ |
| TN::*arcZ* | Mariner transposon insertion in ArcZ (insertion site: 4692989) | This study |
| ∆*arcZ* | Deletion starting from 4692930 to 4693059 | This study |
| ∆*arcZ*::*arcZ* | Complementation of TT01∆*arcZ* by insertion of full length ArcZ sequence | This study |
| ∆*hexA* | TT01 ∆*hexA* (*plu3090*) | ^3^ |
| ∆*hexA*::*hexA*_PacI_UTR | Knockin of hexA with an altered sequence in the 5‘-UTR | This study |
| *hfq*^3xFLAG^ | *∆hfq*::*hfq*^3xFLAG^ | This study |
| ***Xenorhabdus szentirmaii* DSM16338** | WT, Amp^R^ | ^12^ |
| ∆*hfq* | ∆*hfq* (Xsze_00563) | ^13^ |
| ∆*hfq*::pCEP_GXPS | Promotor exchange in front of *gxps* | This study |
| ∆*arcZ* | Deletion starting from 3833948 to 3833859 | This study |
| ∆*arcZ*::*arcZ* | Complementation of XSZ∆*arcZ* by insertion of full length ArcZ sequence | This study |
| ∆*arcZ*::pCEP_GXPS | Promotor exchange in front of *gxps* | This study |
| ∆*hexA* | ∆*hexA* (*Xsze_03702*) | This study |

**Supplementary Table S15**. Oligonucleotides used in this study

| **Name** | **Sequence (5´-3´)** | **Purpose** |
| --- | --- | --- |
| NN191 | TATAACCTCTCCTTAATTTATTGC | Linearization of pSAM_Bt |
| NN192 | AAACAATAGGCCACATGC |  |
| NN193 | TAAATTAAGGAGAGGTTATACTGCGTCTAGCATGCCTA | Amplification of Kan^R^ |
| NN194 | TTGCATGTGGCCTATTGTTTTTAGAAAAACTCATCGAGCATC |  |
| NN276 | ATCGATCCTCTAGAGTCGACCCTGAATGATTTTGATTACGCT | Deletion of *arcZ*: amplification of *upstream* region of *arcZ* (TT01) |
| NN277 | GACGCTGAAAAAAAATAACCCAAAGATAAGATTTTTTGTTACAAGATTC |  |
| NN278 | GGTTATTTTTTTTCAGCGTCC | Deletion of *arcZ*: amplification of *downstream* region of *arcZ* (TT01) |
| NN279 | TCCCGGGAGAGCTCAGATCTCGATGTATTATCAAGTGAAAGGC |  |
| NN281 | GGCAGACTCCTGTAGAACG | Verification primer for TT01∆*arcZ* |
| NN282 | CGCATAAGATAAAAGGTGCTGC |  |
| NN315 | TCCTCTAGAGTCGACCTGCACGAAAAACGATAAAGTTGTGAGC | Deletion of *arcZ*: amplification of *upstream* region of *arcZ* (XSZ DSM) |
| NN316 | GGAACACAAGTACCAACATAGC |  |
| NN317 | TATGTTGGTACTTGTGTTCCCACCCCAACTTCGGTTGG | Deletion of *arcZ*: amplification of *downstream* region of *arcZ* (XSZ DSM) |
| NN318 | GGAATTCCCGGGAGAGCTCAGCAACAGGAACGGGCATTG |  |
| NN329 | CGACACTTCAGCACCAAG | Verification primer for XSZ∆*arcZ* |
| NN330 | GCGAGAGATCAGAAAGGAATTAC |  |
| NN333 | ATCGATCCTCTAGAGTCGACGCACTGCTAAAACGTGTCAG | Knockin of *hexA_Pac*I*_*UTR: amplification of *upstream* region of *arcZ* (TT01) |
| NN334 | TTAGTTAGTAATTAAAATCAAAAAAAAGTGATG |  |
| NN335 | TGATTTTAATTACTAACTAATTAATTAATTTACGTAAGCACTATCAAATTAAATTAACATC | Knockin of *hexA_Pac*I*_*UTR: amplification of *downstream* region of *arcZ* (TT01) |
| NN336 | GGAATTCCCGGGAGAGCTCACCTCCCTCTGAATGTTTTGAAG |  |
| NN346 | ATCGATCCTCTAGAGTCGACGCTTTGCGTGCAGAATATAAC | Deletion of *hexA*: amplification of *upstream* region of *hexA* (XSZ DSM) |
| NN347 | CCTGGTGTTTTTACTTCAGC |  |
| NN348 | GCTGAAGTAAAAACACCAGGGTATTACGTATTTATAGGCTAATGTTTCC | Deletion of *hexA*: amplification of *downstream* region of *hexA* (XSZ DSM) |
| NN349 | GGAATTCCCGGGAGAGCTCACCAATAACACCAAAATGAAAATGC |  |
| NN359 | CAGAATAAGGTGAATTTAGTTGATG | Sequencing of *hexA_Pac*I*_*UTR knockin part 1 |
| NN357 | GATGCAGAAGGTGAACTAAG |  |
| NN358 | CCAGCATACTTTCAATAAACTG | Sequencing of *hexA_Pac*I*_*UTR knockin part 2 |
| NN356 | GGTTCAATAGGTAAAAAAACAGC |  |
| X.sz_Cl2_ORF00346_fw_gib_LP  X.sz_Cl2_ORF00346_rv_gib_LP | TTTGGGCTAACAGGAGGCTAGCATATGAAAGATAGCAAGGTTGCT  TCTGCAGAGCTCGAGCATGCACATGGTATACATGATGTACGCTGG | Promotor exchange in front of *gxpS* |
| X.sz_Cl2_ORF00346_ver_fw_LP  X.sz_Cl2_ORF00346_ver_rv_LP | GGCTGCTGGGAATGACAAT  GATAACGAGCCGGACTACAGC | Verification primer for pCEPKMR_ORF00346 |
| KPO-0009 | CTACGGCGTTTCACTTCTGAGTTC | 5S oligoprobe (*E. coli*) |
| KPO-0092 | CCACACATTATACGAGCCGA | construction of pMH078 (p-*arcZ*) |
| KPO-1397 | GATCCGGTGATTGATTGAGC |  |
| KPO-1702 | ATGCATGTGCTCAGTATCTCTATC | construction of pMH079 (pXG10-*hexA::gfp*) |
| KPO-1703 | GCTAGCGGATCCGCTGG |  |
| KPO-3024 | CAATATGGGGATATCAAAGAAAAGC | RybB oligoprobe (TT01) |
| KPO-3109 | GGACTACACACAGCAATATAGG | RprA oligoprobe (TT01) |
| KPO-3110 | GCTGATTCACTTTTCGTTCCG | GcvB oligoprobe (TT01) |
| KPO-3111 | CTTACCTCTGTACCCTACGC | Spot 42 oligoprobe (TT01 and XSZ) |
| KPO-5989 | GAATACTGCGCCAACACCAG | ArcZ oligoprobe (TT01 and XSZ) |
| KPO-6007 | ACACTACCATCGGCGCTAC | 5S oligoprobe (TT01 and XSZ) |
| KPO-6061 | CTAAAGTAAACACTGGAAGCAATG | RyhB oligoprobe (TT01) |
| KPO-6062 | GCCTGTTGTTTATCTACAGTCAG | GlmZ oligoprobe (TT01) |
| KPO-6130 | AGACAGGGATGGTGTCTATG | CpxQ oligoprobe (XSZ) |
| KPO-6132 | TTAAGAGCCGTGCGCTAAAAG | RyeB (SdsR) oligoprobe (TT01) |
| KPO-6145 | GAGATACTGAGCACATGCATGTCAGAAAACAAAATAACCAAACT | construction of pMH079 (pXG10-*hexA::gfp*) |
| KPO-6146 | CCAGCGGATCCGCTAGCAACAAAAGTTCTTAGCAGATCG |  |
| KPO-6147 | GGCTCGTATAATGTGTGGGTATGATGTACGGAGAATTCC | construction of pMH078 (p-*arcZ*) |
| KPO-6148 | GCTCAATCAATCACCGGATCCAAGAATGGAGAAAGGATACG |  |
| KPO-6149 | CAATATGGGGACATCAAAGAAAAG | RybB oligoprobe (XSZ) |
| KPO-6156 | GTTTTCCCTGCTGTTGGCGCAGTATTCGCG | construction of pMH080 (p-*arcZ**) |
| KPO-6157 | GCGCCAACAGCAGGGAAAACTTTTTACACGC |  |
| KPO-6164 | CGTAAAAACAGCAGGTTAGTTAGTAATTAAAATC | construction of pMH081 (pXG10-*hexA*::gfp*) |
| KPO-6165 | CTAACTAACCTGCTGTTTTTACGTAAGCACTATC |  |
| KPO-6166 | CTGGTGTTGACGGAAATAAGC | sRNA_00243 oligoprobe (TT01) |
| KPO-6169 | GAGGTGGTTCCTAGTCTTAC | CyaR oligoprobe (TT01 and XSZ) |
| KPO-6171 | CGCGAATACTGCGCCAACA | ArcZ* oligoprobe |
| KPO-6303 | GATACACGGTCAGAACACTATC | sRNA_00408 oligoprobe (TT01) |
| KPO-6305 | CCGTAGCGCTGATTGATACC | sRNA_01262 oligoprobe (TT01) |
| AHp292 | TCCTCTAGAGTCGACCTGCACATCAATTTGCTATTGCACC | Introduction of *hfq*^3xFLAG^: amplification of *upstream* product |
| AHp293 | GTAGTCGATATCATGATCTTTATAATCACCGTCATGGTCTTTGTAGTCTTCAGTGCCATCACTTTCC |  |
| AHp294 | TTATAAAGATCATGATATCGACTACAAAGATGACGACGATAAATAGTAAGTTTGAGATCAGTAATAGAATGAAG | Introduction of *hfq*^3xFLAG^: amplification of *downstream* product |
| AHp295 | GGAATTCCCGGGAGAGCTCATATATCCCACCAGTGAAACC |  |
| AHp142 | CGGTTATCGTCAGATGTGG | Verification Primer for Introduction of *hfq*^3xFLAG^ |
| AHp143 | CAACAAAATATTTCGGATGAGG |  |

**Supplementary Table S16**. Compounds analyzed by Target analysis.

| **Compound Name** | ***m/z*** | **Ion** | **Reference** |
| --- | --- | --- | --- |
| Isopropylstilbene (IPS) | 255.1 | [M+H]^+^ | ^14^ |
| Anthraquinone 270a (AQ-270a) | 271.1 | [M+H]^+^ | ^15^ |
| GameXPeptide A (GXP-A) | 586.4 | [M+H]^+^ | ^16^ |
| Photopyrone D (PPY-D) | 295.2 | [M+H]^+^ | ^17^ |
| Phurealipide A (PL-A) | 229.2 | [M+H]^+^ | ^7^ |
| Mevalagmapeptide (MVAP) | 334.8 | [M+2H]^++^ | ^16^ |
| Xenofuranone A (XF-A) | 281.1 | [M+H]^+^ | ^18^ |
| GameXPeptide C (GXP-C) | 552.4 | [M+H]^+^ | ^16^ |
| Protoporphyrin IX (PPIX) | 563.3 | [M+H]^+^ |  |
| Xenoamicin A (XA-A) | 650.9 | [M+2H]^++^ | ^19^ |
| Rhabdopeptide 772 (RXP-772) | 772.4 | [M+H]^+^ | ^20^ |

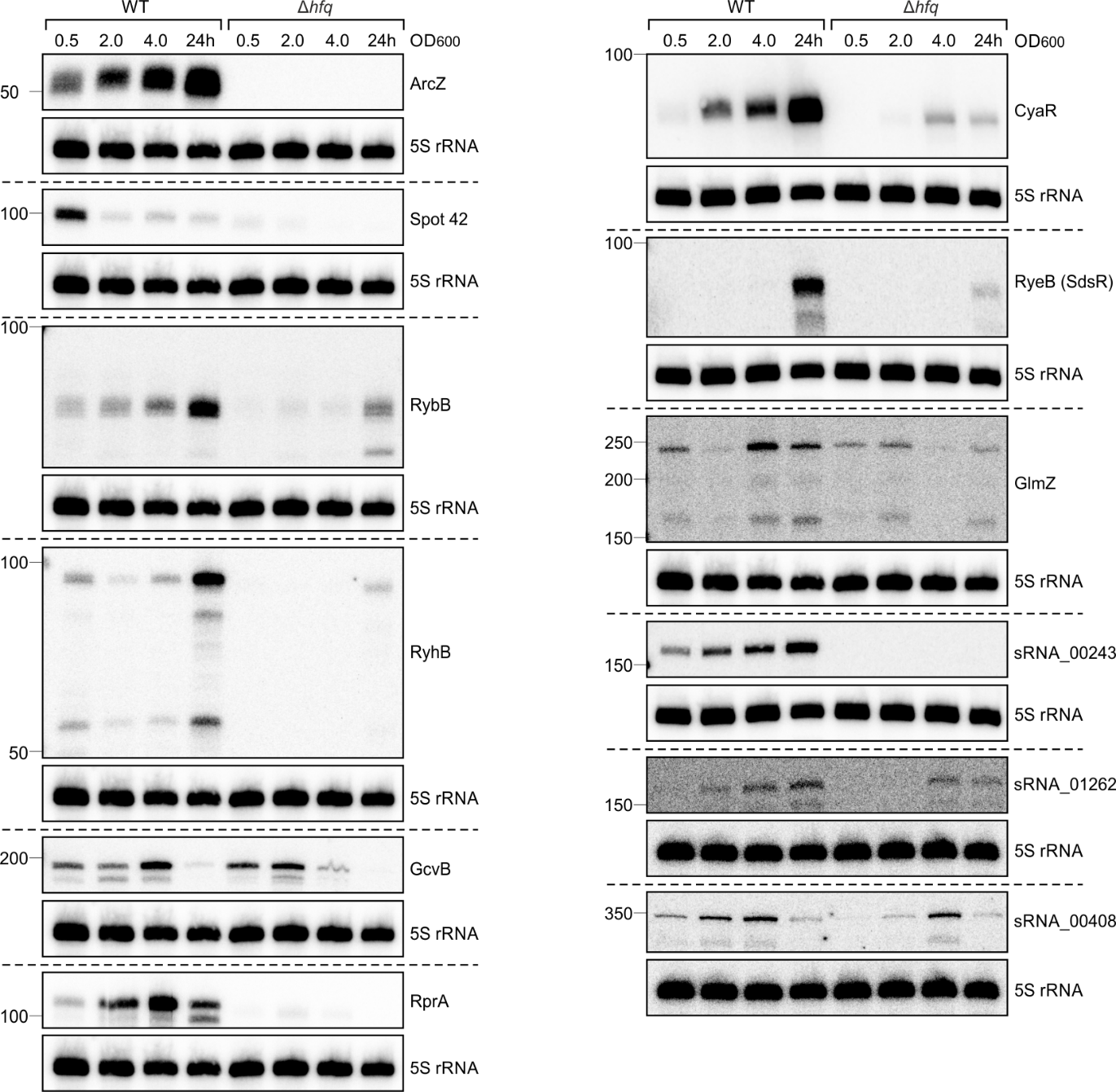

**Supplementary Figure S1.** Expression of various sRNAs in *P. laumondii* at different time points. RNA samples of *P. laumondii* WT and ∆*hfq* strains were taken at three different OD_600_ values (0.5, 2 and 4) and after 24 h of growth. The RNA was loaded on Northern blots and probed for the indicated sRNAs. Probing for 5S rRNA served as loading control.

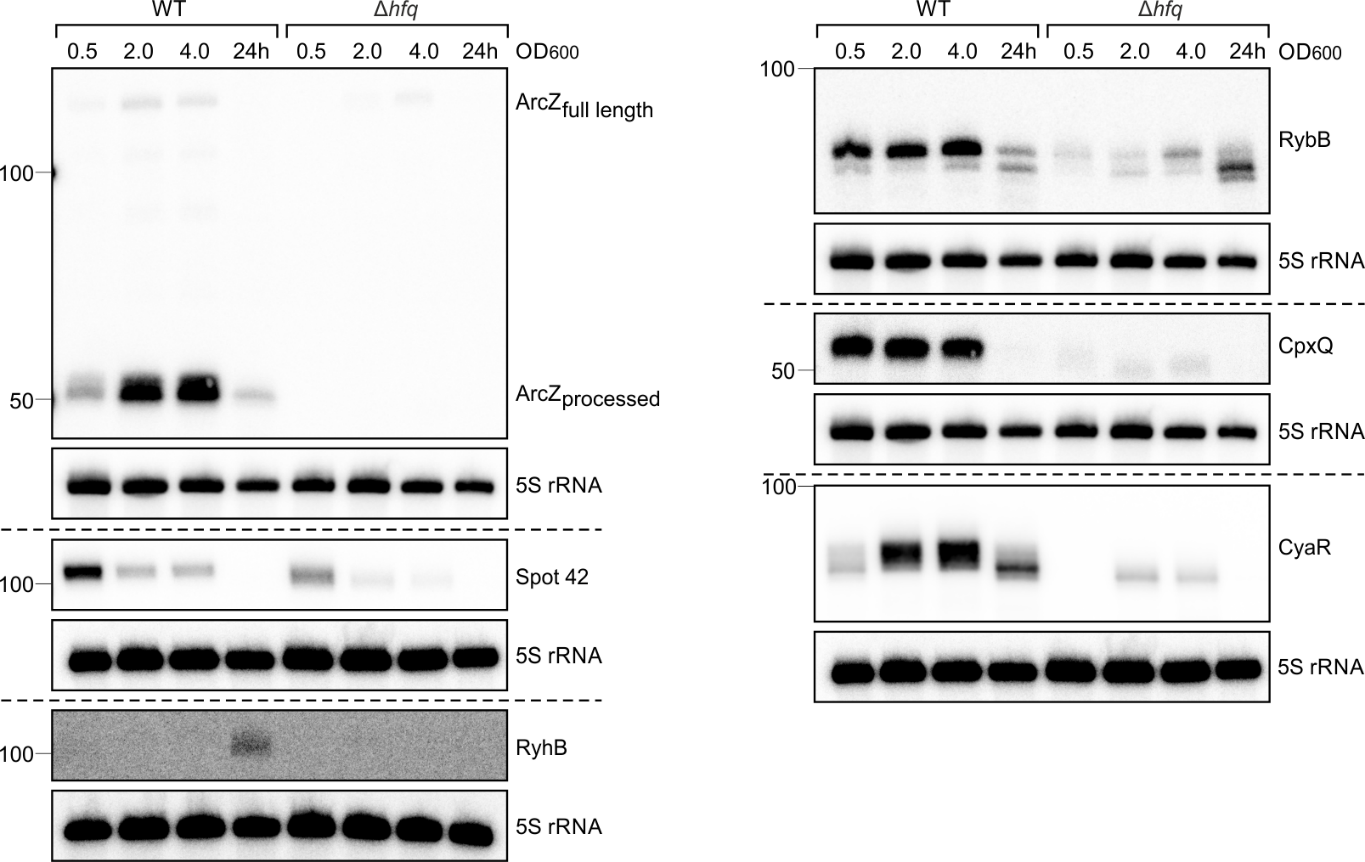

**Supplementary Figure S2.** Expression of various sRNAs in *X. szentirmaii* at different time points. RNA samples of *X. szentirmaii* WT and ∆*hfq* strains were taken at three different OD_600_ values (0.5, 2 and 4) and after 24 h of growth. The RNA was loaded on Northern blots and probed for the respective sRNAs. Probing for 5S ribosomal RNA served as loading control.

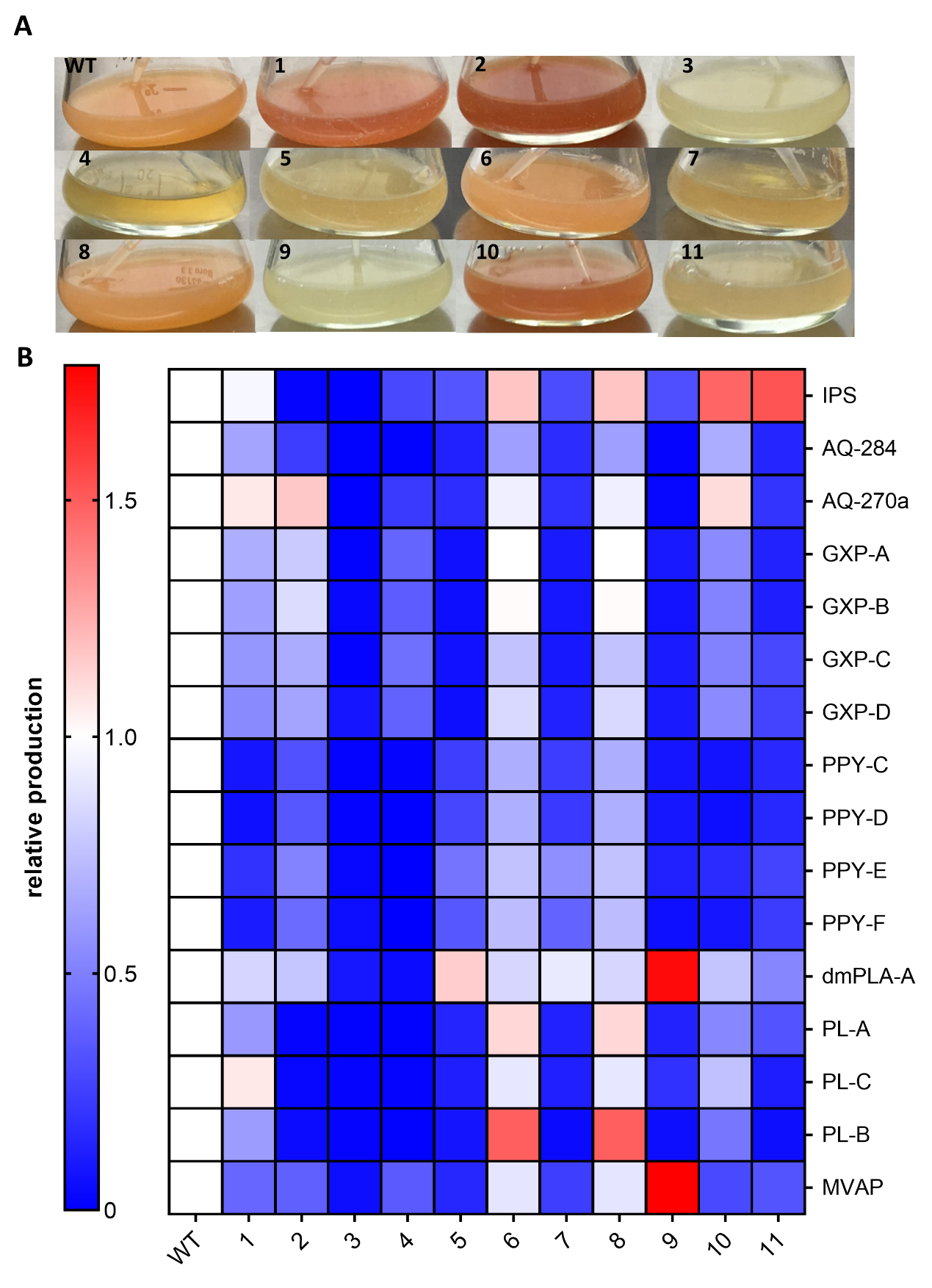

**Supplementary Figure S3.** Phenotype of transposon insertion mutants of *P. laumondii*. **A.** Differences in pigmentation of transposon insertion mutant liquid cultures compared to WT. Depicted are eleven transposon insertion mutants and a WT culture after 3 d of cultivation at 30°C with shaking. **B.** SM- profiles of the transposon insertion mutants. Relative SM production was quantified from duplicates using TargetAnalysis (Bruker) and compared to the WT of *P. laumondii* after 72 h cultivation at 30°C with shaking. Mutant 3 was analysed further and the transposon insertion was identified in the *arcZ* gene.

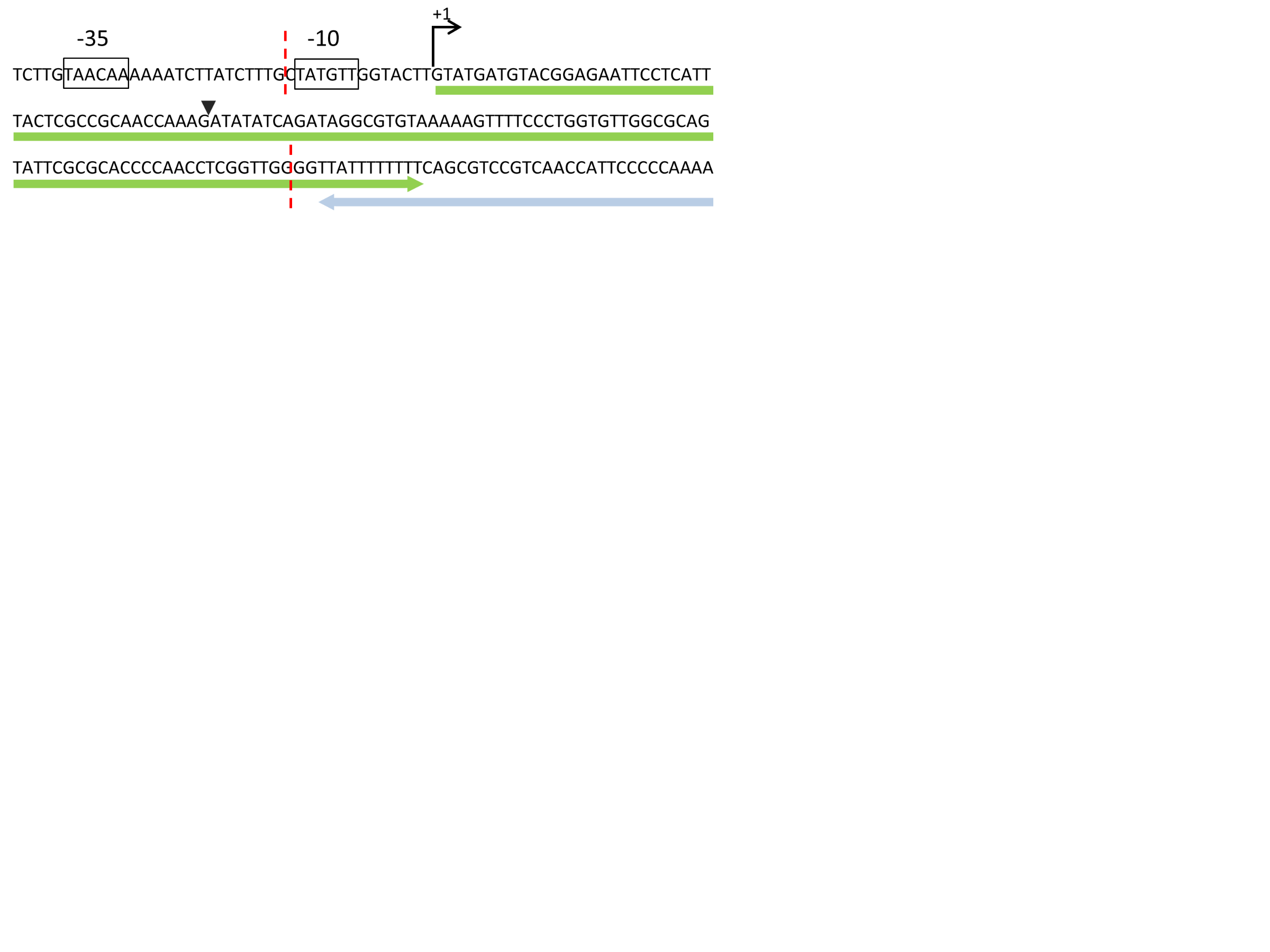

**Supplementary Figure S4.** Sequence of region in *P. laumondii* TTO1 containing predicted *arcZ* sequence (green arrow). The 3’ end of *arcB* (blue arrow) is also shown. Dotted red lines indicate region of *arcZ* that was deleted. Also indicated is the site of insertion from transposon sequencing (inverted black triangle), as well as the -35 and -10 promotor regions and the transcriptional start site (+1).

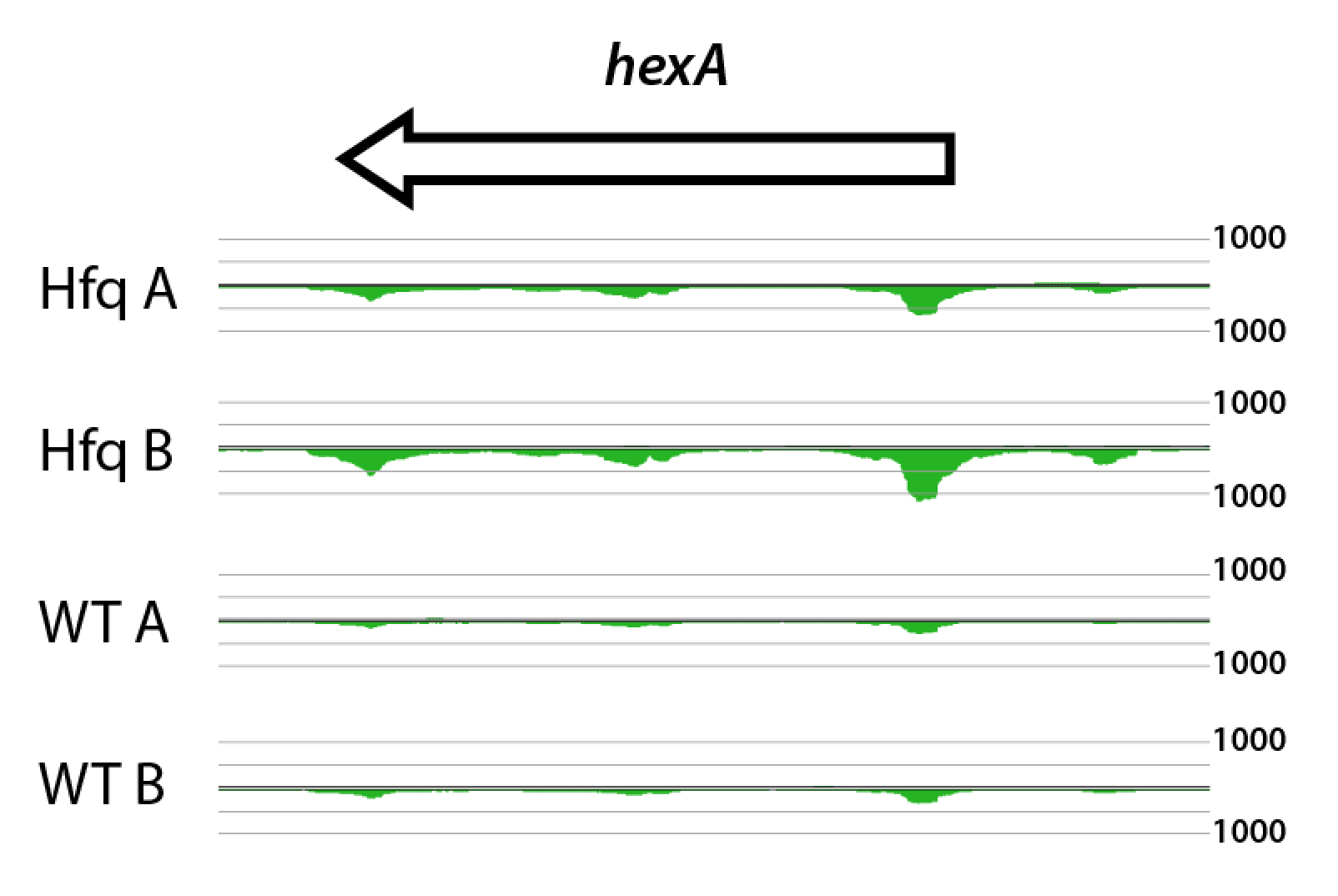

**Supplementary Figure S5.** RIPseq enrichment around the region of *hexA* in Hfq^3xFLAG^ samples (Hfq A & B) and untagged samples (WT A & B). Plots indicate the strand reads map to (bottom = reverse, top = forward). Scale represents perfectly mapped reads. For all enriched regions, see Supplementary Tables S11 and S12.

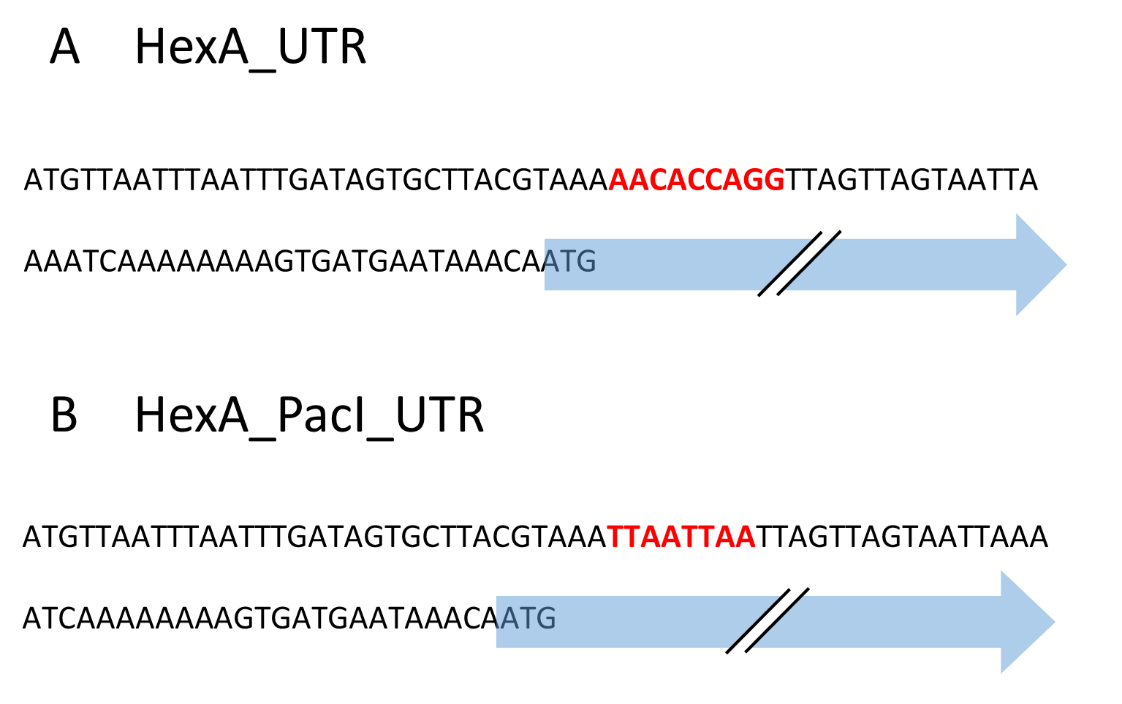

**Supplementary Figure S6. A** 5’-UTR of *hexA* including the predicted ArcZ binding site (red). The arrow indicates the start of the *hexA* coding sequence. **B** The predicted ArcZ binding site (AACACCAGG) was exchanged to a *Pac*I restriction site (TTAATTAA) as shown.

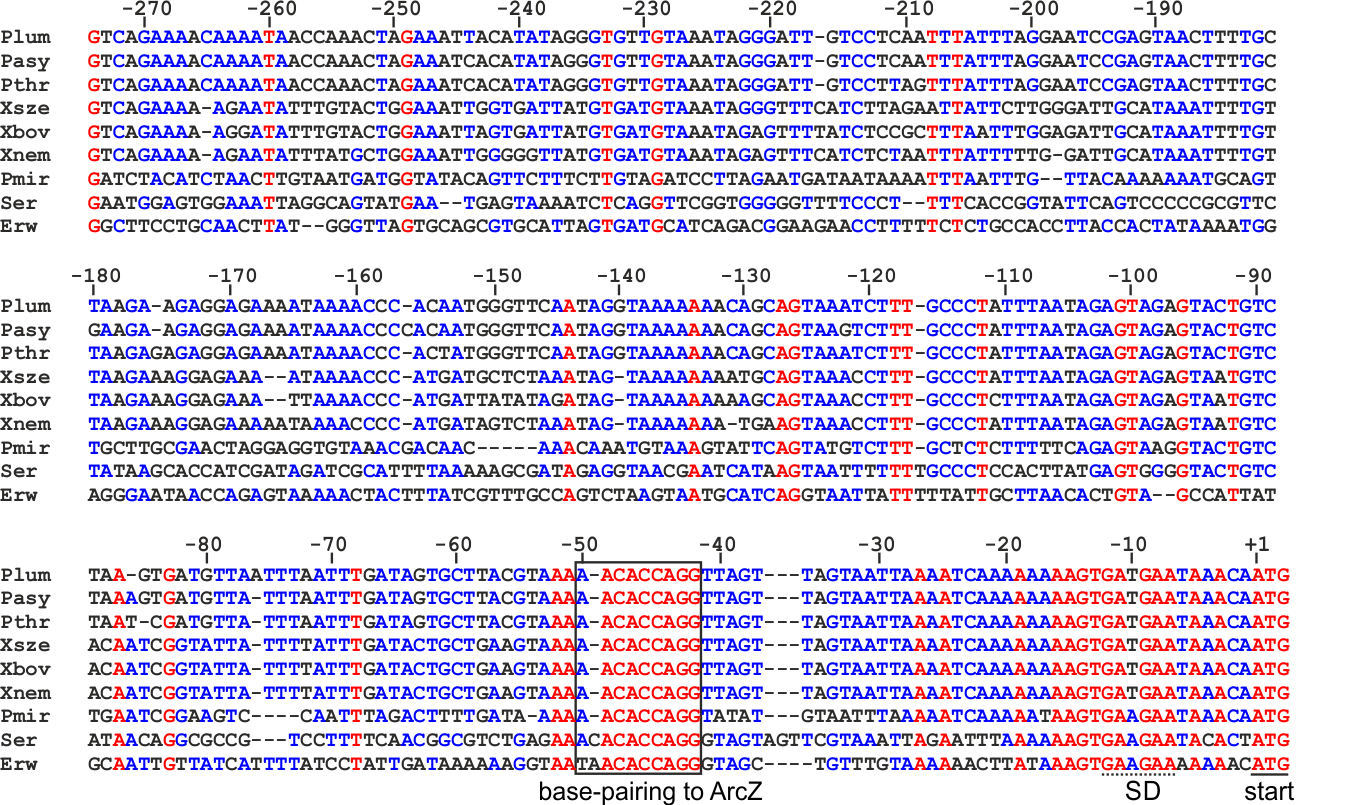

**Supplementary Figure S7**. Alignment of the *hexA* 5’ UTR from *P. laumondii* TT01, *P. asymbiotica*, *P. thracensis*, *X. szentirmaii*, *X. bovienii*, *X. nematophila*, *Proteus mirabilis*, *Serratia marcescens* and *Erwinia* sp. J780, beginning with the transcriptional start site. The sequences were aligned using the Multalign Algorithm^21^. Black box indicates the region of base-pairing to ArcZ. SD sequence and start codon of *hexA* are underlined. Numbers indicate distance to the start codon.

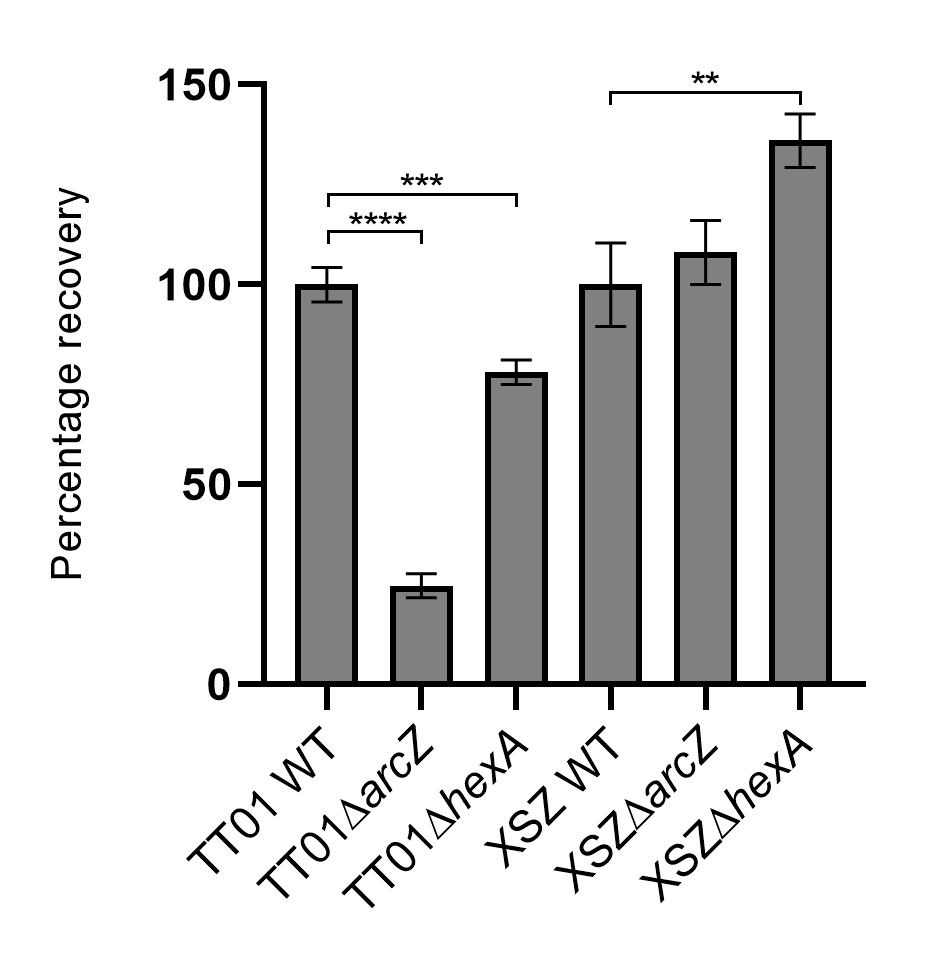

**Supplementary Figure S8.** Infective juvenile development to hermaphrodites with strains of *P. laumondii* and *X. szentirmaii*. Bars represent mean values of 10 individual experiments. Error bars represent the standard error of the mean. Asterisks indicate statistical significance (* p<0.05, ** p<0.005, *** p<0.0005, **** p<0.00005) of relative recovery compared to WT recovery levels.

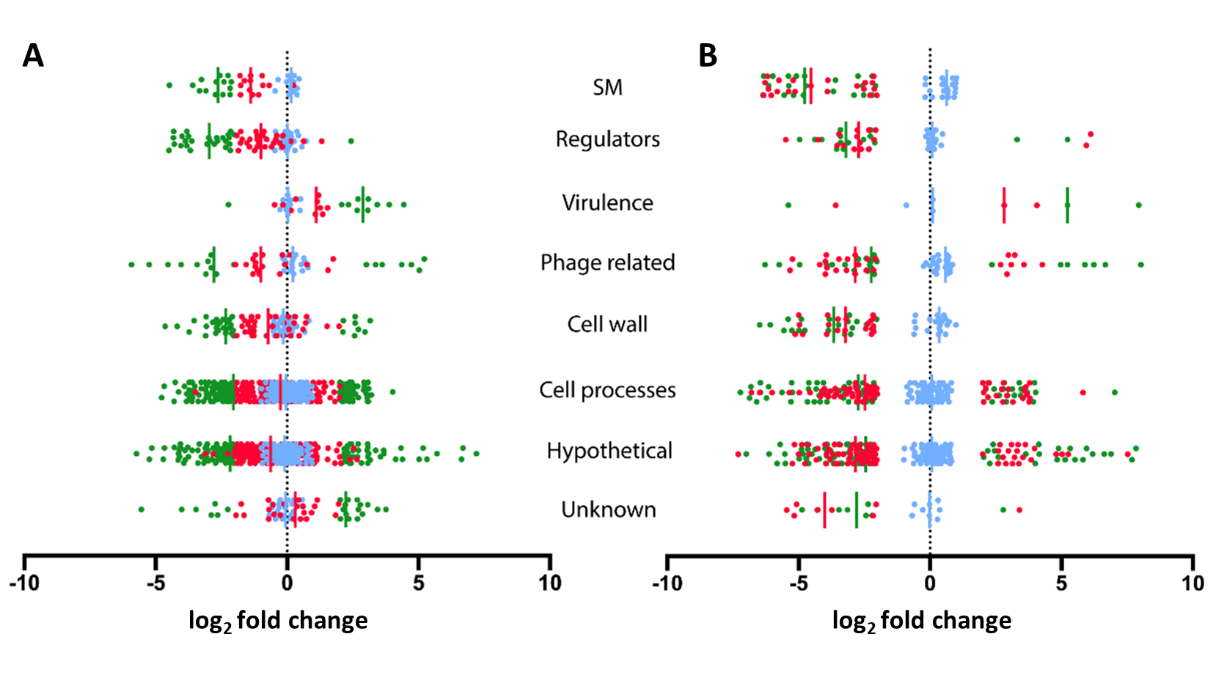

**Supplementary Figure S9A.** Genes that were significantly affected in the ∆*arcZ* strain and not the ∆*hfq* strain or **B** affected in both ∆*arcZ* and ∆*hfq* strains. The coding sequences associated with ∆*arcZ* of *P. laumondii* (green), ∆*hfq* (red) or ∆*arcZ::arcZ* (blue) compared to the WT were grouped into eight different categories: specialized metabolites (SM), regulators, virulence, phage related, cell wall, cell processes, hypothetical and unknown based on their annotations. Vertical lines represent the median for each group. Complete lists of regulated genes for *P. laumondii* mutants can be seen in Supplementary Tables S14-S15.
